# E*x vivo* Model of Functioning Human Lymph Node Reveals Pivotal Role for Innate Lymphoid Cells and Stromal Populations in Response to Vaccine Adjuvant

**DOI:** 10.1101/2024.08.21.608943

**Authors:** Joannah R. Fergusson, Jacqueline H.Y. Siu, Nitya Gupta, Eloise Nee, Sören Reinke, Tamara Ströbel, Ananya Bhalla, Shyami M. Kandage, Thomas Courant, Sarah Hill, Moustafa Attar, Alex Gordon-Weeks, Mark Coles, Calliope A. Dendrou, Anita Milicic

## Abstract

Immunological processes that underpin the administration of therapeutics and vaccines are poorly defined due to a lack of models which faithfully recapitulate human immune responses. Inbred mice lack the diversity inherent to people, while the microanatomical organisation of human tissue is lost in isolated cell suspensions. We describe precision-cut human lymph node (LN) slices as architecturally-preserved, functioning lymphoid tissue model system, and explore early inflammatory responses to a potent vaccine liposomal adjuvant containing a TLR4-agonist and QS21 saponin. Combining scRNA-seq, multiplexed immunofluorescence and secretome analysis, we dissect direct and indirect signalling pathways in both leukocytes and stromal cells to reveal communication networks linking innate and adaptive immunity. Application of molecular inhibitors reveals that secretion of IL-1β, but not IL-18, is TLR4-dependent in human LN. Retaining donor-to-donor immune variation, this *ex vivo* LN model system enables the study of pathways previously difficult to observe in humans, paving the way towards precision medicine.

## INTRODUCTION

Our understanding of the functioning of the human immune system relies largely on animal studies in inbred mice. While enabling both systemic and tissue-specific interrogation of immune responses, animal models lack much of the complexity of humans and are often poorly predictive of the response to immune perturbations such as vaccination or infection^1^. Rational drug and vaccine design requires preclinical models that accurately reflect human immune responses, to aid in the mechanistic characterisation, and accelerate the clinical development, of new preventative and therapeutic interventions^2,3^.

Formerly, modelling human immune responses was restricted to *in vitro* 2D systems such as cell co-cultures or whole blood assays (WBAs)^4^ which offer an over-simplified interpretation of immune interactions, without fully replicating the complex immune environment of the lymphoid tissue where these responses initiate^5^. More recently, ultrasound-guided fine needle aspiration (FNA) of lymph nodes has been utilised to monitor vaccine-induced responses^6–11^ and autoantibody generation^12,13^, enabling temporal sampling of the immune system from within native tissue. However, in addition to a complete loss of spatial information, FNAs predominantly capture T and B lymphocytes with a relative loss of information about rarer innate immune and non-hematopoietic cells^14^ which are critical for induction and maintenance of immunity^15,16^.

Lymph nodes (LNs) are highly organised and structured to enable the cell-cell interactions required to mount an effective immune response. The importance of experimentally recapitulating these interactions to accurately model the *in vivo* responses has long been appreciated^17–19^, and current efforts have turned to designing 3D *in vitro* replicas of human lymphoid tissue, either by combining single-cell suspensions with a matrix scaffold^2^ or through reaggregation of dissociated tissue, as described with tonsil-derived organoids ^20^. While 3D *in vitro* models such as organoids can be informative, the initial tissue disaggregation irretrievably leads to the loss of original, physiological, architecture, and these models often lack critical tissue components such as the stromal compartment or extracellular matrix^15^. An alternative approach is the *ex vivo* culture of non-dissociated tissue block explants, as reported with tonsil^21,22^ and spleen tissue^23,24^, and utilised in studying responses to infection^25^ and vaccination^26^. These explants retain the natural tissue architecture, but given the inherent need to sample only a small subsection of the whole organ may suffer from sampling bias, failing to capture all tissue structures and leading to divergence in responses^23^. Further, tonsil and spleen tissue are less relevant early post-vaccination, with the immune response typically initiated in the draining lymph node.

We sought to profile the early immune events at the single-cell level in the human LN as full-organ cross-sectional live tissue slices, during induction of inflammation by a potent, clinically-relevant vaccine adjuvant. Inflammation is a common pathway in the generation of an effective immune response to vaccination and the induction of long-lived immunity^16^. Vaccine adjuvants are known to induce an effective inflammatory innate immune response, enhancing later adaptive immunity in a successful immunisation^3^, although complete understanding of their mechanism of action has been limited by poor translatability from animal models into clinical efficacy, with even licensed adjuvants lacking fully defined mechanisms^27^. Adjuvant AS01 by Glaxo Smith Kline (GSK) combines a TLR4 agonist with the saponin QS-21 in a liposomal formulation as part of the highly effective licenced vaccines against herpes zoster (Shingrix^®^) and malaria (Mosquirix^TM^), yet our mechanistic insight into its immune mode of action derives mostly from mouse studies^28–31^.

Mechanistic studies are also limited due to the fact that most potent adjuvants remain proprietary. We employed an open-access adjuvant LMQ, similar in composition to AS01, with proven safety and efficacy in animal models^32,33^, to *ex vivo* stimulate tissue slices of live human LNs, and examine the innate immune pathways triggered. From a single-cell transcriptomic map of the early adjuvant-induced events, we observed a pivotal function for innate lymphoid cells (ILCs), particularly NK cells, in translating inflammation into subsequent adaptive immune responses that could lead to enhanced protection. In addition, by stimulating intact tissue, we reveal the key role of resident non-hematopoietic stromal cells, a previously underappreciated cell type in studies of vaccine responses, in the amplification of the inflammatory response within the human LN.

## RESULTS

### 1. Precision-cut human LN slices are transcriptionally and architecturally preserved in culture, reflecting the native human LN

To dissect functional responses of human lymphoid tissue following immune perturbation, we developed an *ex vivo* culture system which allows individual cell populations to be observed in the context of their physiological tissue microenvironment. Healthy, non-inflamed human LNs were obtained through routine elective cholecystectomy, excised from surrounding mesenteric tissue. A method for precision cutting of whole LNs into 300 μm cross-sections was developed (Materials & Methods), with tissue viability retained over short-term culture, allowing a range of analytical assays **(Figure 1A, Suppl. Fig. 1)**. Cell populations present in a whole human LN at baseline remained present and viable during stimulation *ex vivo* **(Figure 1B)**, although a 3.7-fold reduction in the percentage of macrophages was observed in LN slices after culture.

**Figure 1:**
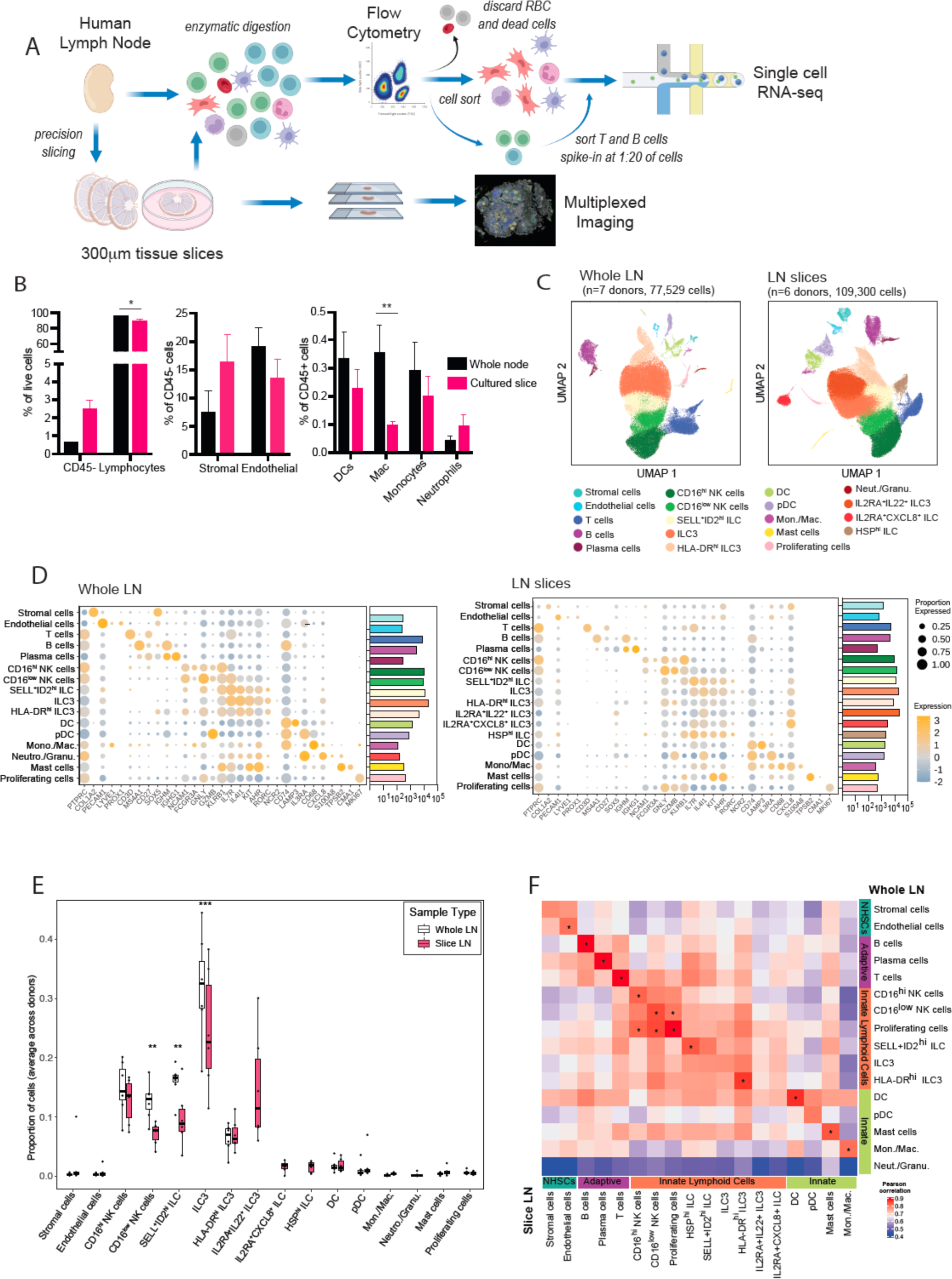
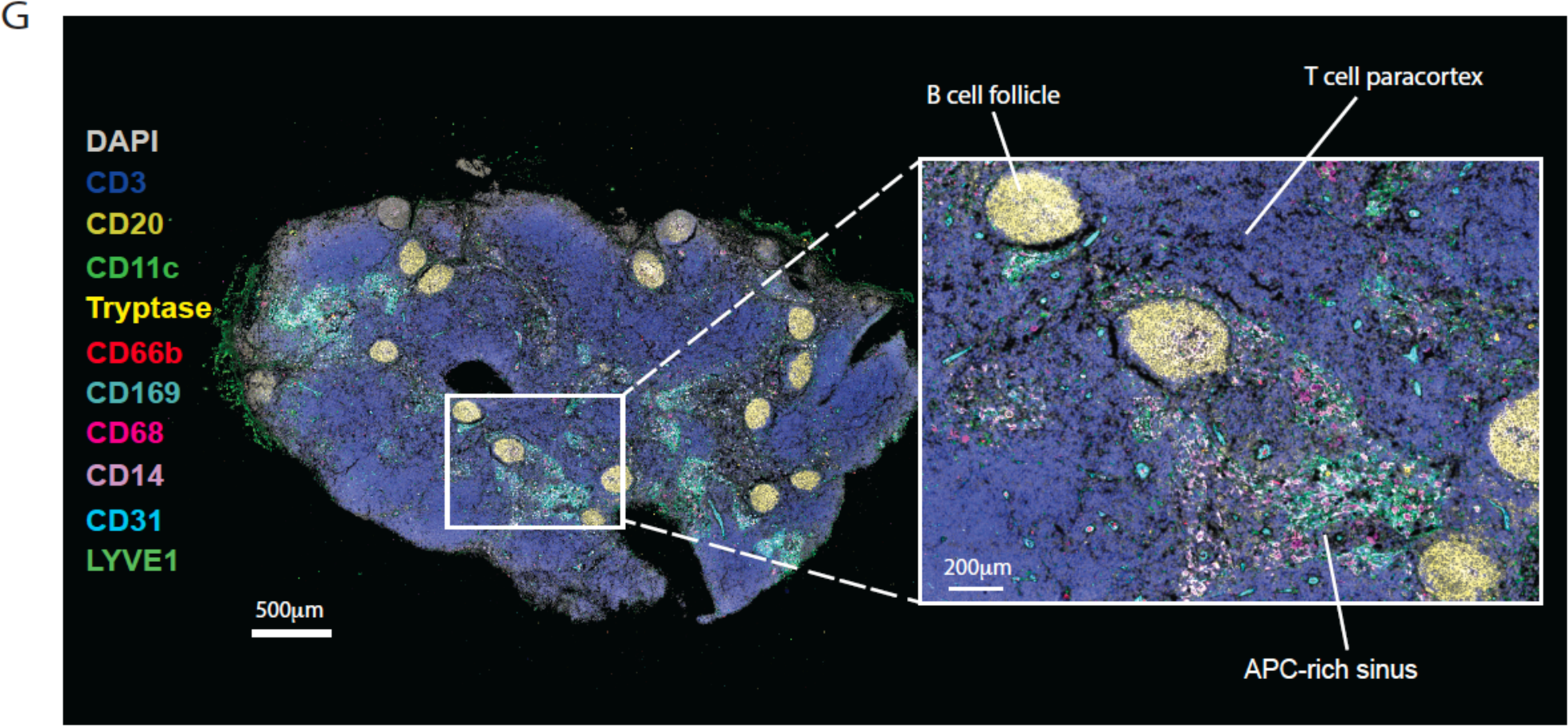
Precision-cut human LN slices are transcriptionally and architecturally preserved in culture to reflect a whole native human LN *ex vivo*. A) Workflow schematic of transcriptomic analysis of whole or sliced human LNs. Live human LNs from individual donors were digested whole or cut into 300μm cross-sections and cultured for 20 h, followed by digestion to single cell suspensions for assessment by flow cytometry. For transcriptomic analysis, in both whole and sliced LNs, dead cells and RBC were discarded and the majority of T and B lymphocytes depleted by cell sorting before being spiked back at a defined ratio. The resulting cell suspensions were profiled by 10X Genomic Chromium single cell RNA sequencing (scRNA-seq). Slices were also formalin-fixed and paraffin-embedded for multiplexed imaging analysis. B) Live cell type distribution from either whole digested human LNs (n=3) or from 300 μm LN slices after 20h in culture (n=3). *p<0.05, **p<0.005 by two-way ANOVA with Sidak’s multiple comparisons test; all other comparisons were not significant. C) UMAP plots of whole human LN from 77,529 cells pooled from 7 individual donor LNs, and human LN slices of 109,300 cells pooled from six individual donors, coloured by cluster and cell type. D) Dot plots showing expression of marker genes for each cell cluster across whole (left) and slice (right) LN datasets. Colour indicates relative log-normalised level of expression across clusters and dot size the proportion of each cluster expressing each gene. Bar plots indicate the absolute cell counts for each cell type. E) Proportion of each cell type (excluding T and B cells) in unstimulated slices (magenta) and whole native LN (black outline), each point represents an individual donor. **p≤0.005, ***p<0.0005 by ANOVA using a linear model between cell types with matched clusters in whole and slice. F) Comparison of gene expression across all cell types between whole LN (vertical) and slice LN (horizontal): heatmap of the Pearson correlation coefficient between matched clusters for the median gene expression of the top 2000 most variable genes with * indicating where r>0.8. G) Multiplexed stained image of a human LN slice section after 20 h in culture with LMQ adjuvant spatially identifying T cells (CD3), B cells (CD20), macrophages (CD68), monocytes (CD14), NK cells (CD56), DC (CD11c), neutrophils (CD66b), mast cells (Tryptase), endothelial (LYVE1, CD31) and stromal (Podoplanin; Pdpn) populations. Inset images show greater magnification of the boxed area, with markers for stromal (top) and leukocyte (bottom) cell types. Scale bars are indicated, and image is representative of two individuals.

To verify that LN slices post-culture are representative of the cell populations and phenotypes present within an intact human LN, we compared their full transcriptomes: LNs were either dissociated directly into single-cell suspensions, or cut into slices and cultured for 20 h in complete RPMI, prior to enzymatic digestion followed by scRNA-seq analysis. Multiple slices (average of 4 per condition) from each donor were pooled after culture to account for differences in the anatomical location of specific cell types. As T and B cells dominate numerically within the LN **(Figure 1B)**, these were sorted separately, and after removing red blood cells (RBC) and dead cells, added (spiked) back at a 1:20 ratio to enable a more balanced representation across all cell populations. Transcriptomic data from 77,529 ‘whole LN’ cells from seven donors, and 109,300 ‘LN slice’ cells from six donors, were separately integrated by donor, followed by unsupervised clustering analysis to identify cell populations across whole or sliced LNs **(Figure 1C)**. Cell clusters in both whole and sliced LNs were annotated using expression levels of canonical gene markers and gene sets **(Figure 1D)**, identifying clusters corresponding to myeloid, stromal and lymphoid populations, including the spiked-in T and B lymphocytes. After T/B cell depletion, the majority of cells within both whole and sliced LNs were innate lymphoid cells (ILCs). This includes NK cells, with two clusters identified based on CD16 transcriptional expression and confirmed at the protein level by flow cytometry and conventional gating strategies ^34^ **(Suppl. Fig. 2)**. In the whole LN, ILCs formed three clusters, all expressing *KIT* (CD117), *IL23R* and *RORC*, associated with an ILC3 phenotype. ILC3s were also the major detected subset at the protein level, with the majority being NKp44**^−^ (Suppl. Fig. 2)**. While an NKp44^+^ population was detected at the protein level, only low levels of its gene locus *NCR2* transcripts were found, consistent with previously described discrepancy between NKp44 transcript and protein expression^35^. Notably, however, *NCR2* transcript levels were similar to those detected in NK cells, and were restricted to the ILC3 cluster of ILCs. Further *NCR* transcripts, *NCR1* (NKp46) and *NCR3* (NKp30), were found at highest levels within the IL2RA^+^SELL^+^ID2^hi^ ILC cluster, which also expressed *GATA3*, and may represent a more naïve ILC subset^35^. Conventional flow-cytometry gating strategies identified ILC1 and ILC2, but these populations were rare and variable between donors and did not form a separate cluster by transcriptional analysis. Three additional ILC clusters were apparent in LN slices, namely IL2RA^+^IL22^+^ ILC3, HSP^hi^ ILCs and IL2RA^+^CXCL8^+^ ILCs, indicative of the plasticity of this cell type and their sensitivity to environmental cues^36–38^ (**Figure 1C**&D). A cluster of neutrophils/granulocytes was detectable in the whole LN but absent from cultured LN slice data, reflecting the difficulty of capturing this cell type for transcriptomic analysis after culture ^39,40^. Cell abundances of matched cell populations beyond ILCs were not significantly different between whole and slice LN datasets, including myeloid cells (DC, monocytes/macrophages, mast cells), pDC, stromal and endothelial cells **(Figure 1E)**. T and B cells were excluded from this analysis due to being depleted and spiked-in at a defined ratio.

To better compare the cell type/states between whole and sliced LNs, the median gene expression of the top-most variable features across datasets was computed for each cluster, and the Pearson correlation coefficient calculated between clusters in whole and slice LN datasets. This confirmed a strong correlation in the gene expression profiles of slice LN cell clusters with matched cell clusters in the whole LN, demonstrating the preservation of cell phenotype *ex vivo* **(Figure 1F)**.

The LN is a highly organised structure and therefore, to faithfully recapitulate its immune pathways, individual cells need to be maintained within their natural microenvironment. Multiplexed imaging of slices confirmed the preservation of LN architecture in culture, including B cell follicles, T cell paracortical regions and sinuses enriched with antigen presenting cells (APCs) **(Figure 1G)**.

### 2. Human LN slices respond to inflammatory perturbation *ex vivo*

Having established the overall preservation of tissue architecture and cell composition, we sought to profile human LN responses as full organ, cross-sectional, live tissue slices during induction of inflammation by a vaccine adjuvant. Adjuvants improve the quality and magnitude of immune responses to vaccine antigen by inducing transient inflammation^4,32,33^. In particular, high potency has been demonstrated for adjuvants that comprise multiple immunostimulatory components, such as AS01^28,31,41,42^. We employed an AS01-like liposome (L)-based adjuvant, containing a synthetic toll-like receptor 4 (TLR4) agonist 3D-6-acyl-PHAD (3D6AP) (M), and QS21 saponin (Q) – abbreviated as LMQ throughout – to study the inflammatory immune response within human LN tissue slices **(Figure 2A)**.

**Figure 2:**
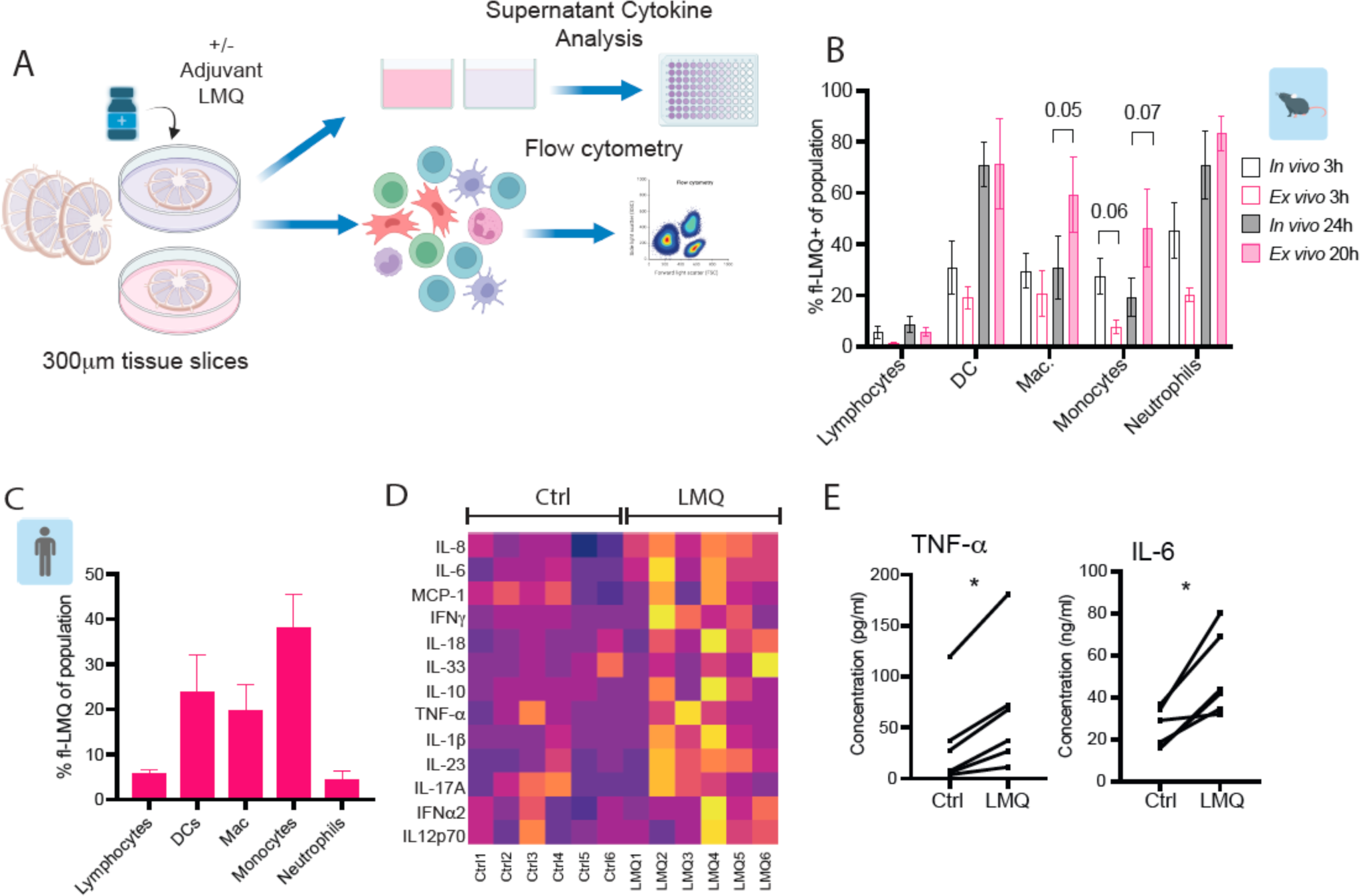
Human LN slices respond to inflammatory perturbation *ex vivo*. A) Workflow schematic of LN slice preparation and analysis. Live human LNs were cut into 300μm cross-sections which were cultured for up to 20 h with or without an immune stimulant (adjuvant LMQ). Following stimulation the slices were digested to single cell suspension for analysis by flow cytometry or scRNAseq, or the supernatants analysed for release of soluble mediators. B) Kinetics and cell type specificity of uptake of a fluorescently-labelled liposomal adjuvant LMQ in the mouse draining LN following intramuscular (i.m.) injection (*in vivo*; n=4) or in cultured LN slices (*ex vivo*; n=3, 1 slice per mouse). Bars indicate mean and error bars standard deviation (SD). All *in vivo* vs *ex vivo* comparisons were non-significant by multiple unpaired t-tests with Welch correction at each timepoint within cell type; p values <0.1 are indicated. C) Proportion of adjuvant positive cell populations within human LN slices after 20 h of *ex vivo* stimulation (n=3). D) Cytokine concentration in slice culture supernatants after 20 h without (Ctrl) or with LMQ adjuvant (LMQ). Heatmap indicates average concentration per donor, averaged from 3-4 slices, and condition scaled by cytokine (row) (n=6 donors). E) TNF-α and IL-6 concentration in supernatants from paired donor conditions without (Ctrl) or with LMQ adjuvant (LMQ) after 20 h. Each point is the average of 3-4 slices, *p<0.05 by Wilcoxon matched-pairs signed rank test (n=6 donors).

To compare adjuvant uptake between LN slices and a native LN *in vivo*, we injected mice intramuscularly (i.m.) with fluorescently-labelled LMQ, and analysed cell type specificity of adjuvant uptake in the draining lymph-node (dLN) by flow cytometry. Equivalent cell populations were positive for the adjuvant at two timepoints, in both LN slices cultured with LMQ and in dLNs following i.m. LMQ injection **(Figure 2B)**, verifying that adjuvant targeting in *ex vivo* LN slices mirrors *in vivo* cell type specificity. Lymphocytes demonstrated the lowest level of adjuvant uptake, with the majority of DCs and neutrophils positive for fluorescent LMQ at later timepoints. No significant difference was observed between *ex vivo* and *in vivo* uptake within cell types, although there was a trend towards lower uptake in monocytes at the earliest timepoint. Therefore, as the proportion of cells positive for the adjuvant increased over time, in subsequent experiments responses were assayed at 20h post-adjuvant stimulation, with the aim of capturing the highest functional impact on each cell type. Transcriptional responses at this timepoint were also found to correlate with adjuvant concentration in culture **(Suppl. Fig. 1)**.

In precision-cut human LN slices cell populations were capable of selectively taking up fluorescently labelled adjuvant *ex vivo* **(Figure 2C)**. In line with results from mouse LNs, a hierarchy was observed between cell types, with the highest uptake among leukocytes seen with myeloid cells, including dendritic cells (DCs), macrophages and monocytes, while lymphocytes displayed little adjuvant uptake.

Cellular take up of adjuvant within human LN slices resulted in proinflammatory cytokine production, with a range of cytokines detected across donors in the culture supernatant **(Figure 2D)**. Assessment of serial LN slices within individual donors further enabled paired comparisons between adjuvant-treated samples and baseline control, pooled by treatment across multiple slices, revealing a significant induction of TNF-α and IL-6 by LMQ **(Figure 2E)**. The same proinflammatory cytokines are detected in the serum and injection site in mice following immunisation with LMQ adjuvant^32^. Crucially, the LN slice model captures the diversity of human responses, with donor variability in the type and levels of cytokines produced. Together, this demonstrates that cellular uptake of adjuvant within LN slices in culture triggers pathways and functional responses that recapitulate the LMQ-induced mediators observed *in vivo*^32^, supporting LN slice culture as a 3D model for *ex vivo* study of adjuvant-induced inflammation in humans.

### 3. LMQ adjuvant directly activates TLR4 and the NLRP3 inflammasome in tissue monocytes/macrophages

To identify which cells are most likely to initiate inflammation by responding directly to the TLR4-agonist in LMQ, we examined whole LN cell populations for the expression of TLR4, its co-receptors, CD14 and LY96, and the associated signalling molecules **(Figure 3A)**. There was little to no expression of these molecules in lymphoid subsets, including innate lymphocytes such as NK cells, while myeloid lineages showed highest expression, in particular monocytes/macrophages which expressed transcripts for TLR4 and both co-receptors. Expression patterns of *TLR4*, *CD14* and *LY96* were similar in the LN slices after stimulation in culture **(Suppl. Fig. 3)**.

**Figure 3:**
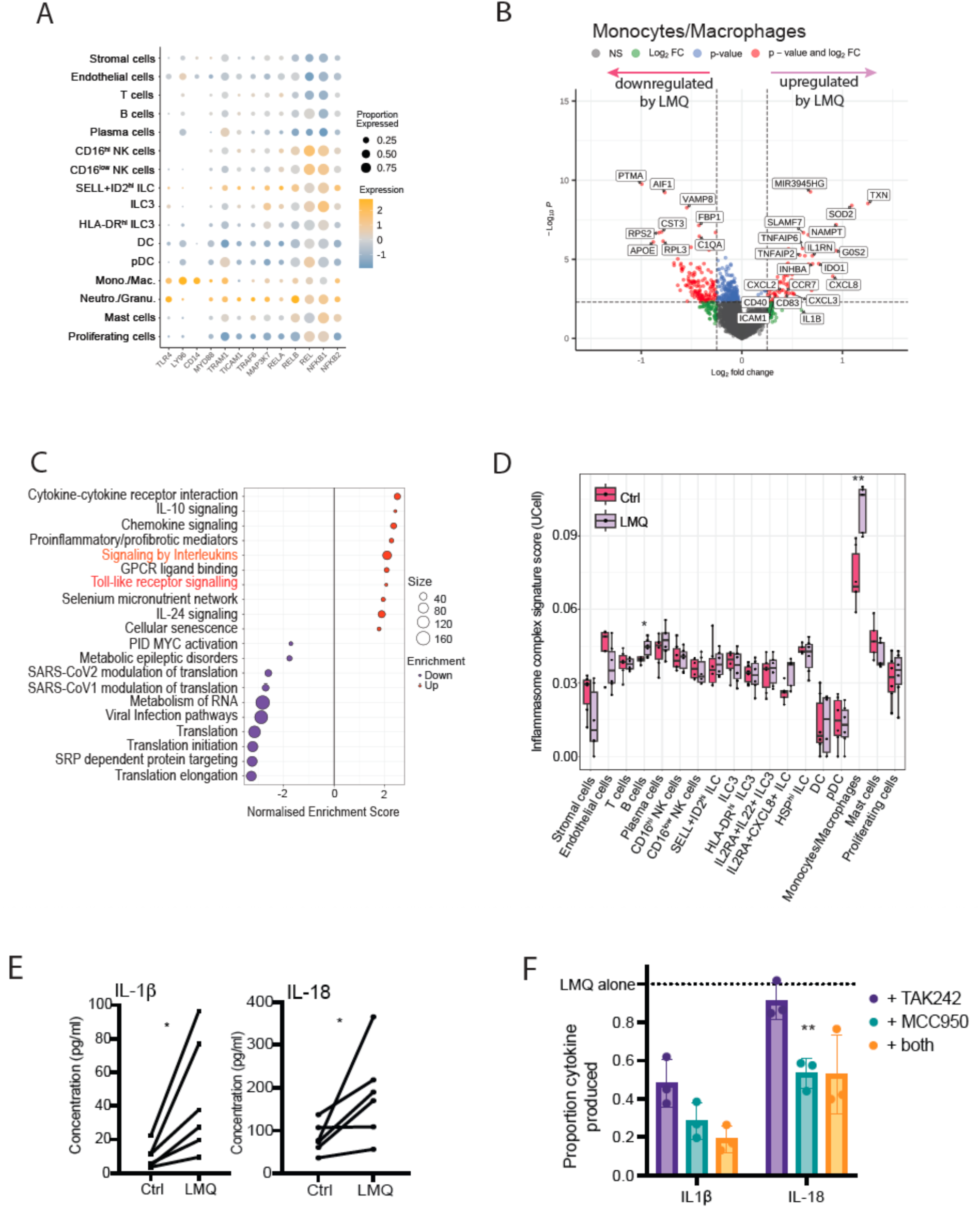
Adjuvant directly activates TLR4 and the NLRP3 inflammasome in tissue monocytes/macrophages. A) Dot plot showing expression of TLR4 signalling components for each cell cluster within whole human LN. Colour indicates relative log-normalised level of expression across clusters and dot size the proportion of each cluster expressing each gene. B) Volcano plot indicating differentially expressed genes between LMQ-stimulated (right: upregulated) and control (left: upregulated) conditions in monocytes/macrophages. Genes with log_2_ fold-change < 0.25 are indicated in green, those with p value <10^-5^ in blue and those fulfilling both criteria in red. C) Enriched canonical pathways among significantly (p<0.05) differentially expressed genes which are up (red)- and down (blue)- regulated in monocytes/macrophages by adjuvant (LMQ) stimulation. D) Gene signature scoring (UCell) of the Canonical Inflammasome Complex Gene Ontology gene set according to cell type and sample type (magenta = Control, lavender = LMQ-treated). The median UCell score for each individual donor are indicated. ** p<0.05 and *p<0.5 by two-way ANOVA with Wilcox test and Benjamini Hochberg correction. E) Concentrations of IL-1β and IL-18 detected in slice culture supernatants after 20 h without (Ctrl) or with LMQ adjuvant (LMQ) from 6 donors, where each point is the average of 3-4 slices *p<0.05 by Wilcoxon matched-pairs signed rank test. F) Relative concentration of IL-1β and IL-18 in slice culture supernatants following 20 h stimulation with adjuvant LMQ in the absence or presence of inhibitors MCC950 (NLRP3 inhibitor), TAK242 (TLR4 inhibitor) or a combination of both. Proportion of each secreted cytokine was calculated relative to LMQ stimulation for each donor (indicated by dotted line at 1.0). (n=3) **p<0.01, *p<0.05 by mixed-effects analysis with Dunnett’s multiple comparisons test to LMQ alone.

The impact of TLR4 signalling was assessed by pairwise differential gene expression analysis within donors, between unstimulated and LMQ-stimulated cell clusters **(Table S2)**. Few differentially upregulated genes were detected in DCs **(Suppl. Fig. 3)**. In contrast, in monocytes/macrophages LMQ stimulation resulted in a transcriptional response including the upregulation of activation markers (*IDO1, CCR7, CD83, CD40*) **(Figure 3B)**. Gene set enrichment analysis (GSEA) of the significantly differentially expressed genes identified TLR signalling among the top enriched pathways in response to LMQ **(Figure 3C; Table S3)**, confirming direct cell activation by the TLR4 agonist. Gene sets related to interleukin signalling were also enriched, suggesting a positive post-activation feedback loop via expression of cytokine mediators.

In mice, LMQ drives inflammation through synergistic TLR4 and QS-21-mediated NLRP3 activation, inducing release of IL-1β and IL-18 ^29,32,43^. We therefore examined the expression of NLRP3 components across LN cell populations. Gene signatures for the canonical inflammasome complex were scored by UCell^44^, and were significantly increased in response to LMQ within the monocytes/macrophages population **(Figure 3D)**, demonstrating the highest median expression across cell types. Interestingly, the inflammasome gene signature was also significantly increased by LMQ in B cells, but to a much lower level. Initiation of the inflammasome pathway results in caspase 1 activation to cleave pro-interleukin 1 and pro-interleukin 18 into their active forms for release, with transcription induced by TLR-mediated priming^45,46^. Both IL-1β and IL-18 cytokines were significantly increased within slice culture supernatants after LMQ treatment **(Figure 3E)**.

To evaluate the relative contributions of TLR4 signalling and NLRP3 activation in LMQ-mediated inflammation, LN slices were pre-treated with molecular inhibitors of these two pathways, TAK242^47^ and MCC950^48^, respectively. MCC950-mediated NLRP3 inhibition reduced the release of both IL-1β and IL-18, while inhibition of TLR4 signalling by TAK242 decreased production of IL-1β but not IL-18 **(Figure 3F)**. This demonstrates that IL-18 is less reliant on TLR4-mediated priming of transcription than IL-1β, consistent with a constitutive expression of IL-18 precursor ^49^, and further indicated by lack of upregulated *IL-18* transcription despite increased protein release **(Suppl. Fig. 3)**. Together, this demonstrates that within the human LN, LMQ initiates inflammation through direct activation of monocytes/macrophages, resulting in the production of inflammatory cytokines including IL-1β.

### 4. Indirect activation of ILCs by vaccine adjuvant leads to downstream signalling to B cells in spatially-preserved LN tissue

Given the maintenance of tissue architecture in LN slices, we investigated the possibility of indirect activation (by inflammatory cytokines) of cell types that lack the ability to directly respond to TLR4 signalling. After depleting T and B cells, ILCs were the major population within the human LN **(Figure 1C)** and with a significant increase in the proportion of IL2RA^+^IL22^+^ ILC3s observed in LMQ-treated slices **(Figure 4A)**. ILC3s can be activated by a combination of IL-1β and IL-23 to release IL-17A, or more particularly IL-22^36,37,50–52^. In addition to elevated IL-1β **(Figure 3E)**, IL-23 was also significantly increased in culture supernatants in response to LMQ **(Figure 4B)**. While IL-17A was not induced at either the transcriptional or protein level **(Suppl. Fig. 4)**, *IL22* transcripts were significantly upregulated within the IL2RA^+^IL22^+^ ILC3 cluster of LMQ treated slices **(Figure 4C)**, and IL-22 concentration was increased in the supernatants of three out of four donors tested **(Figure 4D)**. As LN slice model system retains donor-to-donor variation while enabling paired analyses with multiple readouts **(Figure 2A)**, we assessed whether the supernatant level of IL-22 correlated with the proportion of IL2RA^+^IL22^+^ ILC3s. There was a high correlation (r^2^ = 0.8974) between IL-22 concentration and the proportion of IL2RA^+^IL22^+^ ILC3s in the same donor slices **(Figure 4E)**, providing further evidence that the released IL-22 originates from activated ILC3.

**Figure 4:**
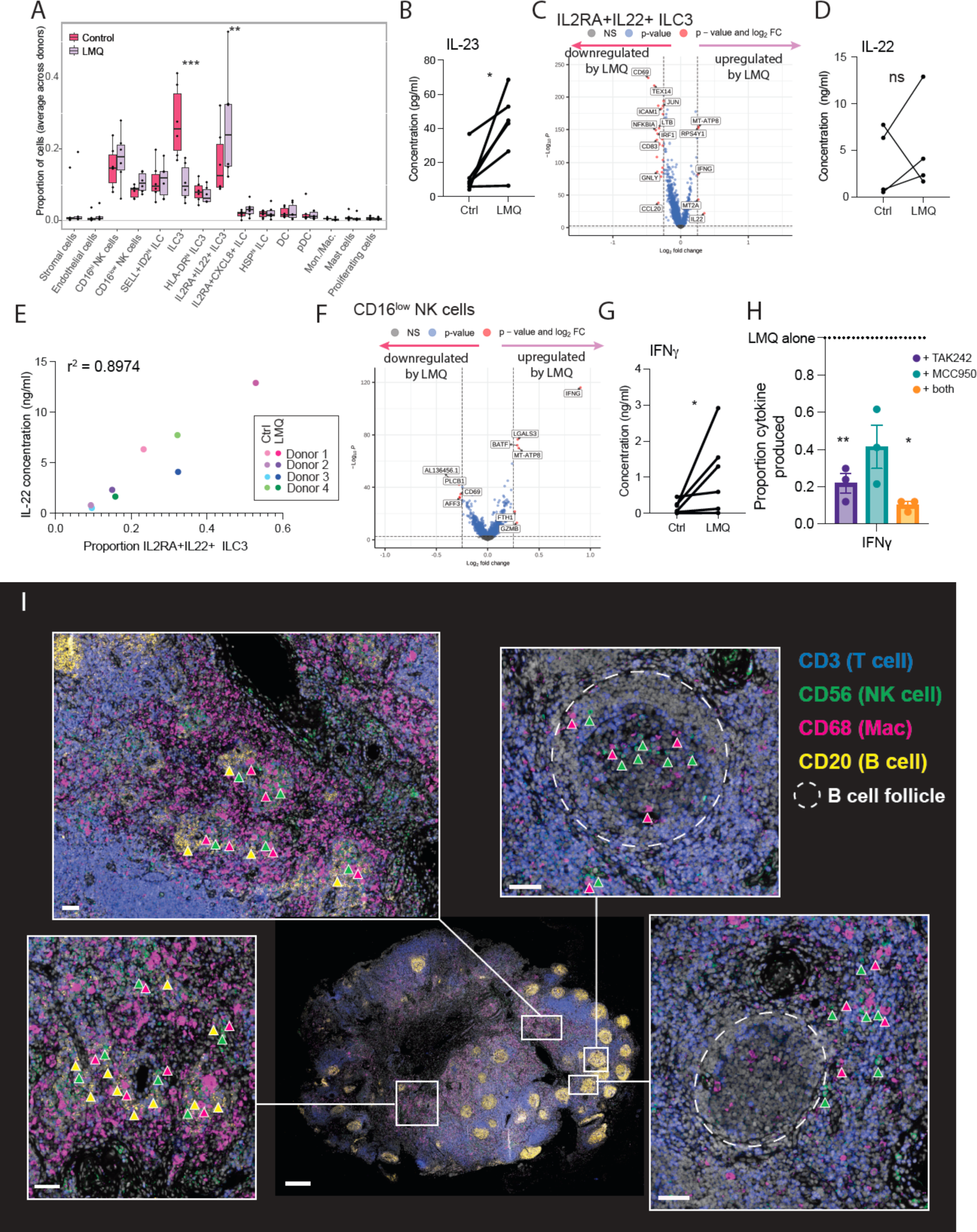
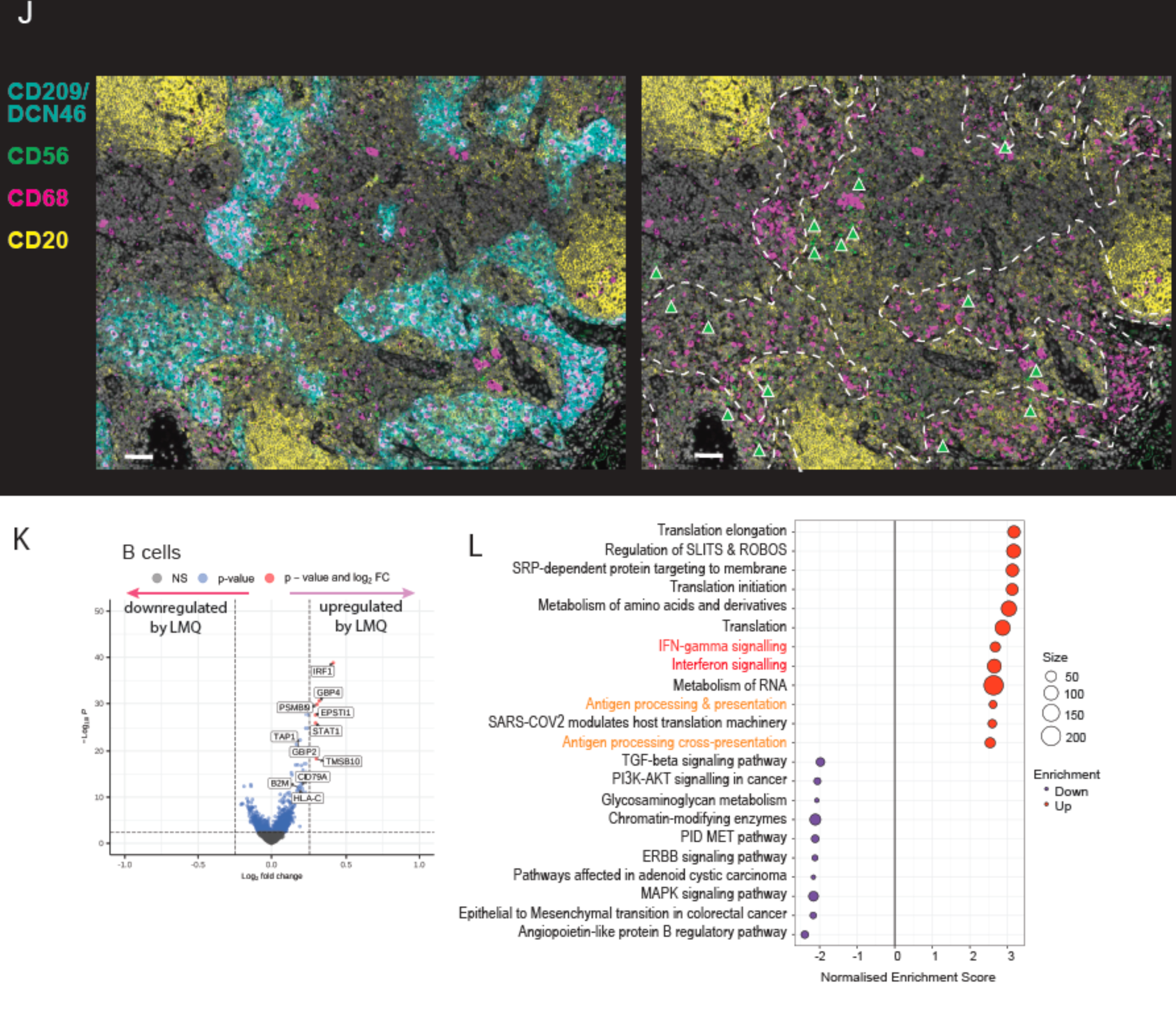
Indirect activation of Innate Lymphoid Cells by adjuvant results in downstream signalling to B cells in spatially-preserved LN tissue. A) Proportion of each cell type in unstimulated (magenta) and LMQ-stimulated (lavender) LN slices, each point represents an individual donor. **p<0.005, ***p<0.0005 ANOVA using a linear model. B) Cytokine concentration of IL-23 (n=6) in slice culture supernatants after 20 h culture without (Ctrl) or with LMQ adjuvant (LMQ) from paired donor conditions where *p<0.05 by Wilcoxon matched-pairs signed rank test. C) Differentially expressed genes in IL2RA+IL22+ ILC3 between LMQ-stimulated (right: upregulated) and control (left: upregulated) conditions. Genes with log_2_ fold-change <0.25 are indicated in green, those with p value <10^-5^ in blue and those fulfilling both criteria in red. D) Cytokine concentration of IL-22 (n=4) in slice culture supernatants after 20 h culture without (Ctrl) or with LMQ adjuvant (LMQ) from paired donor conditions where *p<0.05 by Wilcoxon matched-pairs signed rank test. E) Proportion of IL2RA^+^IL22^+^ ILC3 cells per donor and per condition against IL-22 concentration detected in culture supernatants. Dots are coloured according to donor, and shade according to treatment (light = control, dark = LMQ treated). R^2^ value was calculated by simple linear regression. F) Differentially expressed genes in CD16^low^ NK cells between LMQ-stimulated (right: upregulated) and control (left: upregulated) conditions. Genes with log_2_ fold-change <0.25 are indicated in green, those with p value <10^-5^ in blue and those fulfilling both criteria in red. G) Concentration of IFNγ in slice culture supernatants after 20 h culture without (Ctrl) or with adjuvant LMQ (LMQ) (n=6, each point is the average of 3-4 slices from one donor *p<0.05 by Wilcoxon matched-pairs signed rank test). H) Relative concentration of IFNγ, TNFα and IL-23 in slice culture supernatants following 20 h stimulation with adjuvant LMQ in the absence or presence of inhibitors MCC950 (NLRP3 inhibitor), TAK242 (TLR4 inhibitor) or a combination of both. Proportion of each secreted cytokine was calculated relative to LMQ stimulation for each donor (indicated by dotted line at 1.0). (n=3) **p<0.01, *p<0.05 by mixed-effects analysis with Dunnett’s multiple comparisons test to LMQ alone. I) Multiplexed stained image of whole human LN section to identify location of NK cells (CD3−CD56+, green triangles) in relation to macrophages (CD68, magenta triangles) and B cells (CD20, yellow triangles). Scale bar indicates 400μm. Inset images are zoomed in regions of boxed areas, where scale bars are 50μm. CD20 stain is omitted and B cell follicles indicated with a dashed line in specific images to enable further cell types to be visualised (right). Images are representative of three individuals. J) Multiplexed stained image of whole human LN section to identify location of NK cells (CD56, green triangles) in relation to medullary sinuses (CD209/DCN46 left, dotted line right). Macrophages (CD68) and B cells (CD20) are also shown and scale bars are 50 μm. Images are representative of three individuals. K) Differentially expressed genes in B cells between LMQ-stimulated (right: upregulated) and unstimulated (left: upregulated) conditions. Genes with log_2_ fold-change <0.25 are indicated in green, those with p value <10^-5^ in blue and those fulfilling both criteria in red. L) Enriched canonical pathways among significantly (p<0.05) differentially expressed genes which are up- (red) and down- (blue) regulated in B cells in response to LMQ stimulation.

In addition to IL-1β, LMQ also induced IL-18 release **(Figure 3E)**, which in combination with other innate cytokines such as IL-12, activates NK cells to produce IFNγ^53,54^. One of the most clearly induced genes overall was *IFNG* within CD16^low^ NK cell subset **(Figure 4F)**, consistent with results observed in mice after vaccination with AS01-adjuvanted vaccines^28^. *IFNG* transcription was low in T cells, which generally showed little transcriptional change, and the contribution from other ILC subsets was also low **(Suppl. Fig. 4, Table S2)**, indicating NK cells as the primary source of IFNγ. IL-12 was not detected in culture supernatants **(Suppl. Fig. 4)**, implying that IL-18 synergises with an alternate innate cytokine to induce *IFNG* transcription. Upregulated transcription was accompanied by elevated IFNγ in slice culture supernatants in response to LMQ **(Figure 4G)**. Release of IFNγ was also reduced by pre-treatment of slices with both TLR4 and NLRP3 inhibitors **(Figure 4H)**, despite the lack of TLR4 receptors or inflammasome upregulation in NK cells (**Figure 3**A&D), further supporting their indirect activation by LMQ.

To determine NK cell location and identify the neighbouring cells that provide a potential source of activating cytokines, we turned to multiplexed imaging of the LN slices. CD3^−^CD56^+^ NK cells were identified both within the lymph node medulla and, as rarer events, within or around B cell follicles **(Figure 4I)**. Using antibody clone DCN46, commonly used to detect CD209, which identifies lymphatic endothelial cells and sinus APCs^55^, NK cells were found both within and directly adjacent to the medullary sinuses **(Figure 4J)**. In all cases, NK cells were found next to CD68^+^ cells, suggesting macrophages as the likely source of NK cell-activating cytokines in response to LMQ.

Interestingly, NK cells outside of the B cell-rich follicles were still found in proximity to CD20^+^ B cells **(Figure 4I)**. B cell transcriptional profiling revealed an interferon signalling response to LMQ, including significant upregulation of *IRF1* and *STAT1* **(Figure 4J)**, and enrichment of interferon signalling pathways **(Figure 4K; Table S3)**. As type I interferons were not upregulated in the culture supernatants **(Suppl. Fig. 4)**, this suggests that B lymphocytes responded to paracrine IFNγ from NK cells. In addition, B cells displayed upregulated expression of gene sets associated with protein translation and antigen processing and presentation **(Figure 4K)**, which have also been associated with higher antibody response to vaccination^56^. This reveals a potential mechanism by which adjuvant signals indirectly to the adaptive arm of the immune system, through direct stimulation of myeloid populations and via NK cell-produced IFNγ, ultimately enhancing humoral immunity.

### 5. LN stromal populations are primed both directly and indirectly by vaccine adjuvant to mediate inflammatory cell recruitment

The effect of vaccine adjuvants on resident non-haematopoietic LN stromal populations is rarely studied yet these cells are critical in shaping the immune milieu^15^. Stromal cell populations are also architecturally maintained within LN slices **(Figure 5A**; **Figure 1B)** and interestingly, both mesenchymal stromal and endothelial cells (ECs) showed uptake of fluorescently-labelled adjuvant both in slices **(Figure 5B)** and *in vivo* in mouse dLN following i.m. injection **(Suppl. Fig. 5)**. Exploring LMQ-induced gene expression changes in non-hematopoietic cell populations revealed a strong transcriptional response in ECs **(Figure 5C)**, including gene set enrichment of the TLR signalling pathway: *JUN*, *TAB2*, and *MAPK8* **(Figure 5D; Table S3)**. Although expression levels were low, ECs expressed transcripts for *TLR4*, *LY96*, *CD14* and all downstream signalling molecules in whole human LNs **(Figure 2E)**. Together, this suggests that LMQ directly signals to ECs and induces transcriptional activation through TLR4 stimulation. The IL-1 signalling pathway was also enriched, indicating a potential role for IL-1β in a further indirect activation of ECs.

**Figure 5:**
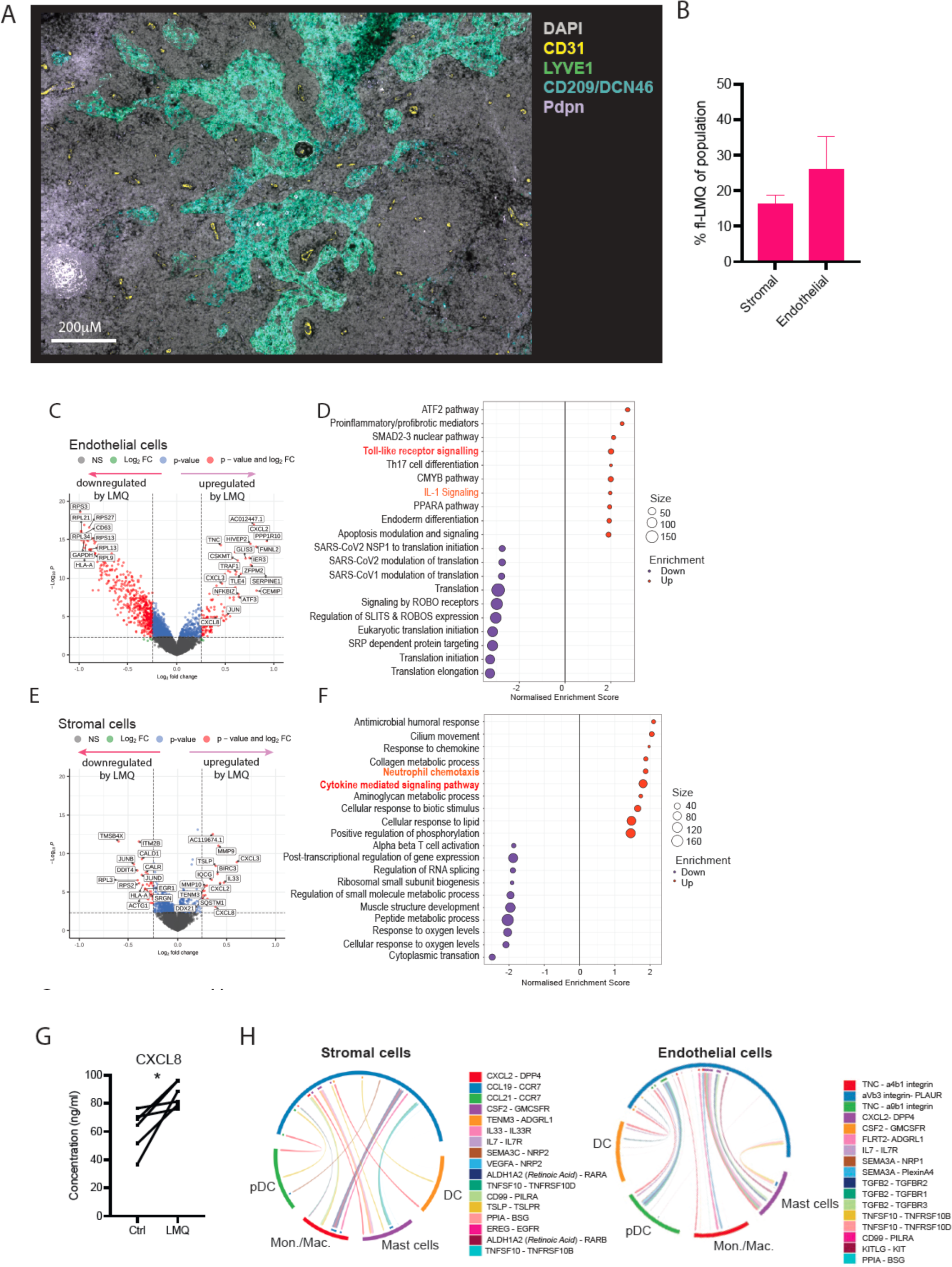
LN stromal populations are primed both directly and indirectly by vaccine adjuvant to mediate inflammatory cell recruitment. A) Multiplexed stained image of a human LN slice section after 20 h in culture with LMQ adjuvant spatially identifying endothelial (LYVE1, CD31, CD209) and stromal (Podoplanin; Pdpn) populations. Scale bars are indicated. Image is representative of results from two donors. B) Proportion of adjuvant positive cell populations within CD45-populations of human LN slices after 20 h of culture (n=3). Stromal cells were defined as CD45−Pdpn+CD31- and endothelial cells as CD45−CD31+ C) Differentially expressed genes in endothelial cells between LMQ-stimulated (right: upregulated) and control (left: upregulated) LN slices. Genes with log_2_ fold-change < 0.25 are indicated in green, those with p value <10^-5^ in blue and those fulfilling both criteria in red. D) Enriched canonical gene sets among significantly (p<0.05) differentially expressed genes which are up- (red) and down- (blue) regulated in endothelial cells following adjuvant (LMQ) stimulation. E) Differentially expressed genes in stromal cells between LMQ-stimulated (right: upregulated) and control (left: upregulated) LN slices. Genes with log_2_ fold-change < 0.25 are indicated in green, those with p value <10^-5^ in blue and those fulfilling both criteria in red. F) Enriched gene ontology biological processes gene sets among significantly (p<0.05) differentially expressed genes which are up- (red) and down- (blue) regulated in stromal cells following adjuvant (LMQ) stimulation. G) Concentration of CXCL8 in slice culture supernatants after 20 h without (Ctrl) or with adjuvant LMQ (LMQ) (n=6, each point is the average of 3-4 slices from one donor *p<0.05 by Wilcoxon matched-pairs signed rank test). H) Relevant interactions upregulated in response to adjuvant stimulation between stromal (left) and endothelial (right) cells and indicated innate cell types as calculated by CellPhoneDB. Interactions were deemed relevant if expressed in more than 10% of cells and identified as differentially expressed upon LMQ stimulation (see Methods). Arrows in the chord diagram indicate the direction of the interaction, with coloured bars indicating the sender cell type and the colour of the bar the receiver cell type.

Examining LMQ-induced gene expression changes within the mesenchymal stromal cell cluster **(Figure 5E)** highlighted enrichment in cytokine-mediated rather than Toll-like receptor signalling pathways **(Figure 5F; Table S3)**. This suggests that the stromal response to LMQ activation is mainly due to indirect, cytokine-mediated, activation rather than direct signalling. Clear upregulation in the transcription of cytokines such as *IL33* and *TSLP*, and of some chemokines was observed, particularly those associated with neutrophil chemotaxis such as *CXLC2 and CXCL3* **(Figure 5E&F)**. Upregulation of these chemokines was also observed in ECs **(Figure 4C)**, indicating this as a common response to LMQ in human LN non-hematopoietic cells. Neutrophil chemoattractant CXCL8 was also significantly increased in culture supernatants following LMQ stimulation **(Figure 4E)** and *CXCL8* was transcriptionally upregulated in multiple cell clusters, including ILCs **(Suppl. Fig. 5)**. *CXCL8*, *CXCL2* and *CXCL3* were all transcriptionally upregulated within monocytes/macrophages **(Figure 3B)**. Indeed, neutrophils were rapidly recruited to the dLN in Balb/c mice after i.m. injection of LMQ, while monocytes and DCs accumulated more steadily over 16h **(Suppl. Fig. 5)**, likely reflecting an early production of neutrophil chemo-attractants by multiple cell types. As mice lack a direct CXCL8 orthologue, the CXCL8 production detected here is unique to human LNs, and CXCL1 production has been reported in mice post vaccination with LMQ^31,32^.

Inferred cell-cell communication, by CellPhoneDB analysis^57^, indicated significant crosstalk in response to LMQ between both stromal cells and ECs and innate cells **(Figure 5H, Table S4)**, across multiple cytokine/chemokine-receptor interactions, including *CXCL2*, *IL33*, *TSLP* and *TNFSF10* (TRAIL). Together, this reveals how stromal cells and ECs are primed by adjuvant to recruit and support innate cell mediators of inflammation for a downstream amplification of the immune stimulus, and illustrates their key role as orchestrators of the response to vaccination in human lymphoid tissue.

## DISCUSSION

Rational design of immune modulators and vaccines relies on a thorough understanding of both their mechanism of action and the immune processes they aim to perturb. To date, much of this knowledge comes from studies in animal models. Yet, due to the relative lack of translation of efficacy to humans, the advantages of studying human responses early during the development pipeline are beginning to be appreciated^2,27^. Evolutionary divergence means that certain immune pathways are not present in the rodent animal models, and thus their contribution cannot be ascertained^1^. On the other hand, genetic and environmental divergence in people results in a huge variation in responses^56,58^ which cannot be captured by the genetically and environmentally controlled laboratory animals. Here we present a precision-cut LN slice model in which the response to an inflammatory stimulus can be studied in a relevant human tissue using a variety of readouts, and reveal mechanistic insights into the triggered processes while retaining the donor-to-donor variation inherent to humans.

Historically, immune mechanistic studies in people have largely been performed in 2D culture systems, generally based on cell populations from peripheral blood and/or cell lines^2,4^. While these can provide specific insights into cell activation and signalling responses, and direct comparisons across formulations^4^, people ages^59^ or between continents^60^, they both oversimplify the immune cell types involved and overlook the role played by the microenvironment. Hence, efforts have turned to lymphoid 3D culture systems, such as organoids or ‘organ-on-a-chip’ models, which have been shown to recapitulate germinal-centre type features *in vitro*^20^. B and T cells from peripheral blood aggregate into lymphoid follicle structures within a matrix gel when cultured under constant flow perfusion in a microfluidic device^61^, while in tonsil organoids single-cell suspensions spontaneously reaggregate on permeable membranes and can be maintained over several days, enabling tracking of recall immune responses to stimulants such as influenza antigens^20,62^. Tonsil organoids have been used to demonstrate the early importance of innate immune cells, such as myeloid and pDC, in antibody responses to influenza vaccine.^20^ However, these approaches seek to reconstruct the original tissue architecture *in vitro*, and hence lose much of the precisely orchestrated anatomy of the native lymphoid tissue. Indeed, the spatial location of both innate leukocytes^16^ and stromal cells^15^ plays a critical role in driving downstream immune responses.

To study native human LN responses, while preserving the tissue microanatomy, we developed a model system for precision cutting and subsequent culture of LN slices. Culturing tonsil explants has previously been used to study vaccines^26^ and infectious agents such as Human Immunodeficiency Virus (HIV)^22,25,63^, although this method relies on using sections or biopsies which are likely to select for specific tissue regions. Precision cutting has been successfully applied in other organs^23,24,64–66^ and species^67^, and the small size of human LNs allows to obtain full-organ cross-sectional tissue slices. Here we demonstrate that 300 μm thick cross-sections of human LNs retain stromal niches and leukocyte populations over short-term culture, enabling direct study of early immune responses. The viability of the tissue over longer time *ex vivo* remains to be fully determined, or indeed the impact of media perfusion which would be important in preventing hypoxia for extended periods of culture^68^.

Employing a clinically-relevant vaccine adjuvant LMQ, a liposomal formulation with a TLR4 agonist and QS21 saponin, we studied the early induction of inflammation within human lymphoid tissue. In mice, following intramuscular injection, LMQ activates the NLRP3 inflammasome and induces pro-inflammatory cytokines in the dLN^32^. After *ex vivo* stimulation with LMQ, our LN slice model demonstrated cell-type specificity in adjuvant uptake equivalent to the mouse dLN, along with inflammasome activation and secretion of NLRP3-dependent proinflammatory cytokines IL-1β and IL-18. The results align with the findings of NLRP3 activation with AS01 adjuvant^43^, which is similar in composition to LMQ. Through the application of specific small molecule inhibitors TAK242 and MCC950 in the slice culture, we further demonstrate that both TLR4 and inflammasome activation are required for IL-1β but not IL-18 production in the human LN.

This adjuvant composition is associated with induction of Th1-type immunity through IFN-γ-mediated cytokine production^28,32^, with IFNγ signalling required for T cell functionality in mice^28^. Correspondingly, in human LN slices LMQ triggered significant production of IFNγ by NK cells, following direct activation of monocytes/macrophages and production of IL-18 that led to NK cell activation^28,29^. The role of the antigen was not investigated here, and may account for the lack of T cell activation observed, aside of the majority T cell depletion. Similarly, we did not observe significant activation of DCs, described to be essential for the AS01-driven T cell responses in mice^29,31^. However, AS01 demonstrates preferential activation of monocytes over myeloid DCs in human peripheral blood in the absence of antigen^69^. A DC maturation signature (upregulation of *CCR7, CD83, IDO1, ICAM1*) in response to LMQ was evident within the monocytes/macrophages cluster and could indicate a role for monocyte-derived DCs in T cell activation within the LN. Indeed, LMQ has been shown to induce the differentiation of human monocytes into DCs, with a Th1-polarising and CD8+ priming potential in 2D *in vitro* models^4^. Further studies are required to determine the impact of the antigen on the early inflammatory processes induced by vaccine adjuvant.

Responses to IFNγ signalling were detected in B cells, which were observed in proximity to both NK cells, shown to produce IFNγ, and the monocytes/macrophages capable of activating them. Identification of this sequential cell-cell communication was only made possible through preserving architecturally intact LN tissue. Direct inflammasome activation by LMQ in B cells may also play a part, as some NLRP3 component upregulation was observed here, although at much lower levels than in monocytes/macrophages. B cell transcriptomes were also enriched for antigen processing and presentation pathways, important in obtaining T cell help and enhancing antibody responses^56,70^. It is interesting to note that NK cells were found in and near medullary sinuses where antibody-secreting B cells reside^71^, facilitating the paracrine signalling through IFNγ. Indeed, IFNγ is upregulated in the serum of people vaccinated with AS01-adjuvanted vaccines^28^, with IFNγ signalling associated with improved adaptive responses^41^, and antibodies considered critical in mediating protection^42^.

This study also highlights both direct and indirect activation of resident LN stromal cells, which are generally neglected in the context of responses to vaccination. Clear upregulation of chemokines such as *CXCL2* and *CXCL3* in mesenchymal stromal and ECs indicate a pivotal role in cell recruitment and amplification of the inflammatory response within the LN. While in mice TLR4 expression within stromal cells was suggested to be dispensable for optimal adjuvant responses, with noted stromal contribution in the maturation of DCs, the impact on cell recruitment was not assessed^72^. Indeed, in stromal cells we observed upregulated expression of *TSLP*, a cytokine shown to activate DCs^73^, as well as *IL33*, an alarmin critical for anti-viral T cell responses^74^. It should be noted that *ex vivo* tissue slices represent a static system, with cell populations uniformly exposed to the adjuvant, rather than temporally through drainage through LN sinuses and conduits^75^. Nonetheless, we show that equivalent cell populations take up the adjuvant in LN slices and in whole dLN after i.m. injection in mice, including structural stromal populations that also demonstrate a significant uptake *in vivo*. Similarly, the effects of cell recruitment cannot be observed, limiting study to resident cell types. Soluble antigen, including adjuvant^29^, has been shown to reach the dLN within minutes, whereas influx of migrating cells takes several hours to many days^31,76,77^ implicating the resident cells as drivers of the earliest immune responses. One exception to this are neutrophils, with antigen-bearing neutrophils found in the dLN within the first few hours post vaccination^31,78^. Although neutrophils were detected in the LN at baseline, they were not retained in the slice cultures and their contribution could not be examined.

Previous analyses of LN transcriptomic responses including LN FNA studies^6–11,61^, have been dominated by lymphocytes, or alternatively involved sorting specific cell populations^79,80^. Here, depletion of T/B lymphocytes enabled us to observe a previously unappreciated role for ILC populations beyond NK cells in the adjuvant-driven inflammatory response, and in particular, activation of ILC3 to produce IL-22, typically expressed by lymphoid tissue inducer (LTi) cells during lymphoid organogenesis.^81^ LTi cells are also involved in the restoration of lymph node integrity following infection in adults^82^. IL-22 has been found to promote chemokine expression in the stromal cells of tertiary lymphoid organs important in regulating B cell responses^83^, offering another mechanism by which adjuvant may be impacting humoral adaptive immunity. In the human LN slices studied, ILC3s were the major non-T/B cell population. Anatomical differences in the distribution of ILC subtypes have been described^34^, and the ILC3 phenotype prevalence within the cystic LN here may relate to its abdominal location, although ILC3s are also the predominant ILC type in hepatic^52^, mesenteric and lung-draining lymph nodes^34^, and thought to be tissue-resident^84^. In mice, both resident and recruited ILC1s are an early rapid source of IFNγ following AS01-adjuvanted vaccination, yet are significantly outnumbered by IFNγ+ NK cells, and both the vaccine platform and the immunisation site are suggested to impact the ILC2 activation phenotype^85,86^. It remains to be investigated in humans whether other ILC subtypes, particularly those in lymph nodes draining the standard vaccination sites, may also be activated by adjuvant. In addition to resident ILC populations, further heterogeneity in cell types, both resident and migratory, exists between LNs from different anatomical sites, and draining distinct organs or regions^87,88^. Further work is required to systematically assess heterogeneity in the immune pathways of LNs of different location and immune history, with human LN slice culture offering a platform through which this could be achieved.

The levels of IL-22 induced by LMQ in LN slice supernatants showed donor variability, but by paired transcriptomic analysis were found to be proportional to the IL2RA^+^IL22^+^ ILC3 population. The induction of inflammatory cytokines was also variable between donors, with some displaying high baseline cytokine levels. Preserving the individual immune variation is important in faithfully recapitulating human responses, such as differences between demographics (e.g. with age) or inherently among individuals. Some donors are described to have a “naturally adjuvanted state”^56^, which may be due to past or current infections and cannot be easily replicated in animal models. Importantly, tissue slices enable the assessment of baseline and treatment within the same individual, accounting for intrinsic differences and infection history, facilitating the study of the impact of such factors on immune responses to drugs or vaccine modalities, across human populations.

The *ex vivo* human LN slice approach described here offers a versatile platform for studying mode of action and early cell signalling responses, and could be applied to a variety of compounds, from novel small molecule drugs to a range of immunostimulants and existing therapeutics. Cumulatively, evaluating responses to immune or inflammatory perturbation on a per donor basis, and building our understanding of the principles and nuances of the underlying mechanisms, could pave the way for the rational design of vaccines and drugs towards achieving precision medicine.

## ACKNOWLEDGMENTS

This study was funded by the Chan Zuckerberg Initiative. J.R.F. was supported by the Chan Zuckerberg Initiative. S.R. was funded in part by the Deutsche Forschungsgemeinschaft (DFG, German Research Foundation, 495054088). J.S. was supported by Chan Zuckerberg Initiative and the Wellcome Trust (226938/Z/23/Z). S.H. was supported by the Wellcome Trust (HMR05310). A. M. was supported by the Chan Zuckerberg Initiative, John Fell Fund (University of Oxford), MRC Confidence in Concept Awards, and the “Adjuvants for Global Health” grant (INV001759) funded by the Bill and Melinda Gates Foundation. C.A.D. was supported by the Chan Zuckerberg Initiative, Wellcome Trust and Royal Society (204290/Z/16/Z), UK Medical Research Council (MR/T030410/1), Rosetrees Trust (R35579/AA002/M85-F2), Cartography Consortium funding from Janssen Biotech, Inc and the NIHR Oxford Biomedical Research Centre, Inflammation Across Tissues and Cell and Gene Therapy Themes. Adjuvant used in this study was provided by the Vaccine Formulation Institute. Sample collection was supported by NIHR Biomedical Research Centre, Oxford. The views expressed are those of the authors and not necessarily those of the NHS, the NIHR or the Department of Health. Thank you to the TGU Investigators for their help in sample collection, and to Paul Klenerman and Nick Provine for help with obtaining ethical approval. Thank you also to Ryan Waters for provision of pig lymph nodes, and to the Kennedy Institute facility staff including Jonathan Webber (Kennedy Flow Facility), Ida Parisi, Rhiannon Cook (Kennedy Histology Facility) and Dylan Windell (Kennedy Digital Pathology Omics Core). Thank you to Sam Pledger and Rowie Borst for lab management. We thank Romain Guyon for helpful discussions and statistical help, Sarah Davidson for assists throughout the project, Floriane Auderset and Marcelle van Mechelen for critical review of the manuscript and Jelena Bezbradica Mirkovic for help with project discussions and interpretation of inflammasome results. Figure schematics were made with BioRender.

## MATERIALS AND METHODS

### RESOURCE AVAILABILITY

#### Lead contact

Further information and requests for resources and reagents should be directed to and will be fulfilled by the lead contact, Anita Milicic.

#### Materials availability

This study did not generate new unique reagents. VFI adjuvant LMQ can be made available through contact via the VFI website (https://www.vaccineformulationinstitute.org/).

#### Data code and availability

Single-cell RNA-seq data will be deposited at CellxGene and made publicly available through CellxGene Discover upon publication. All original code is publicly available on Github (https://github.com/DendrouLab/LymphNodeCultureSlice_2024). Microscopy data and flow cytometry data reported in this paper will be shared by the lead contact upon request. Any additional information required to reanalyse the data reported in this paper is available from the lead contact upon request.

### EXPERIMENTAL MODEL AND STUDY PARTICIPANT DETAILS

#### Human Samples

Cystic lymph nodes (LNs) were obtained from patients undergoing routine cholecystectomy without acute inflammatory pathology or suspicion of malignancy. LNs were resected following gallbladder excision and processed directly after surgery. Donor demographics are provided in Table S1. Written informed consent was obtained from each patient and samples were collected under REC 21/YH/0206. The samples were kept anonymous and handled according to the ethical guidelines set by NHS Health Research Authority.

#### Animals

For all experiments, mice were maintained at the Wellcome Centre for Human Genetics or the Kennedy Institute of Rheumatology, University of Oxford. Animals were house under Specific Pathogen Free (SPF) conditions and in accordance with the recommendations of the UK Animals (Scientific Procedures) Act 1986 and ARRIVE guidelines. Protocols were approved by the University of Oxford Animal Care and Ethical Review Committee for use under Project License PP0984913 granted by the UK Home Office.

For *in vivo* adjuvant uptake experiments 7-10 week old female inbred BALB/C mice (Envigo, UK) were immunised intramuscularly with a total volume of 50μl containing 25μl of fluorescent LMQ in the tibialis muscle under light isoflurane anaesthesia. Draining inguinal lymph nodes were harvested at the indicated time points and digested in RPMI containing 0.8mg/ml Dispase, 0.2mg/ml Collagenase-P and 0.1mg/ml DNase, with mixing at 37°C for 3x 20 mins incubations. Cells were collected in cold PBS supplemented with 2% FCS and 2mM EDTA and filtered through a 100μM cell strainer, before being stained for flow cytometric analysis.

For *ex vivo* adjuvant uptake experiments, inguinal LNs were obtained and cut into 300μM thick slices and cultured with fluorescent LMQ for the indicated timepoints. After culture, slices were digested and stained for flow cytometric analysis as above.

## METHOD DETAILS

### Reagents used are listed in Table S5

#### Adjuvant

LMQ adjuvant was manufactured at the Vaccine Formulation Institute as described previously^33^. Briefly, liposomes composed of 1,2-dioleoyl-*sn*-glycero-3-phosphocholine (DOPC, Merck-Avanti, USA) and cholesterol were combined with QS-21 (Desert King International, USA) and synthetic TLR4 ligand 3D-6-acyl-PHAD (3D6AP) (Merck-Avanti, USA). LMQ was used *ex vivo* at a dose of 12.5μl contained 2.5μg of QS-21 and 1μg of 3D6AP. Fluorescent adjuvant was made by the addition of 1,1’ – Dioctadecyl-3,3,3’3’-Tetramethylindodicarbocyanine Perchlorate (DiD’/DiIC_18_(5)) oil (ThermoFisher, cat no. D307).

#### Precision lymph node slicing and culture

LNs were dissected from surrounding fat and underwent a 1 second wash in 0.01% Digitonin (Thermo Scientific), followed by 5-7 seconds in PBS, to remove residual lipid droplets as previously described^89^, prior to embedding in 2.5% Agarose. Slices were cut from embedded LNs using a Compresstome (Precisionary Instruments), following manufacturer’s guidelines and with a stainless-steel blade set at a speed of 1-2 and oscillation of 6. Surrounding agarose was removed from cut slices and slices were rested in 1ml of complete media (RPMI + 10% FCS + 1x Glutamine + 50U/ml Pen/Strep + 50uM ϕ3-mercaptoethanol + 1mM sodium pyruvate + 1x non-essential amino acids + 20mM HEPES) for 1hr at 37°C before further culture in 1ml of fresh complete media. Each slice was cultured separately. For stimulations, 12.5μl of LMQ (Vaccine Formulation Institute) was added and slices cultured for 20 hrs. For inhibition experiments, slices were pre-treated with 1μM MCC950 (PZ0280, Sigma Aldrich) and/or 5μM TAK242 (614316, Merck) for 1hr before addition of LMQ.

#### Cell and library preparation for single-cell mRNA sequencing

LNs were mechanically disrupted with scissors before being enzymatically digested in RPMI containing 0.8mg/ml Dispase, 0.2mg/ml Collagenase-P and 0.1mg/ml DNase, with mixing. This was performed at 37°C for 3 x 20 mins incubations for whole LN preparations, and 2 x 15 mins for slice preparations. Slices from treatment groups were pooled. After digestion cells were collected in cold PBS supplemented with 2% FCS and 2mM EDTA and filtered through a 100μM cell strainer. Cells were stained with live/dead near infra-red (nIR), anti-CD3, anti-CD20, anti-CD235a, and 7AAD added directly before sorting with the SONY SH800S cell sorter. Dead cells and red blood cells were removed by sorting (CD235a−7AAD−Live/Dead −), while T (CD3+CD20−) and B (CD3−CD20+) cells were sorted separately and spiked back at ratio of 1:20 of sorted non-T/B (CD3−CD20−) cells. Antibody details are listed in Table S6.

Freshly sorted cells were counted and processed according to 10X Genomics guidelines. Single-cell RNA-sequencing libraries were prepared according to the manufacturer’s instructions using Chromium Single Cell 3’ Library and Gel Bead Kit v2 (10X Genomics) and Chromium Single Cell A Chip Kit (10X Genomics) or single-cell RNA sequencing by the GEX 3’ Chromium kit and a maximum of 20,000 cells were loaded per channel.

#### Sequencing and Data Processing

Expression libraries were sequenced by the Oxford Genomics Centre or Novogene on the NovaSeq6000 S4. Each sample was sequenced to an average depth of approximately 36,000 reads per cell.

Cell Ranger v7.0.0 count, with include intron parameter set to TRUE, was used to align reads to the GRCh38-2020-A human transcriptome and generate feature-barcode matrices from the Chromium single-cell RNA-sequencing output. Panpipes was used to perform quality control, batch correction, and clustering pipelines^90^. Quality control for high-quality single cells include removing cells expressing fewer than log1p(6) genes, and cells with mitochondria gene count percentage greater than30.

#### Highly variable gene selection, dimension reduction, clustering, and annotation

UMI counts were normalised by the total number of UMIs per cell, and converted to transcripts-per-10000, and then log-normalised^91^. Top 2000 highly variable genes were selected using vst method from Seurat^92^ and implemented in Scanpy^91^. T cell receptor and immunoglobulin genes were removed from highly variable gene list. Data was then scaled prior to PCA. Harmony was used for batch correction by donor^93^. Leiden clustering was applied to determine cell populations^94^. Further sub-clustering on myeloid populations was required to derive the following populations: Monocytes/Macrophages, DC, Neutrophils/Granulocytes.

#### Comparison of transcriptomes whole vs slice

Median gene expression was determined for each gene across cell clusters in whole and control slice samples. Genes were then filtered to the union of the top 2000 most variable features across both datasets. The Pearson correlation coefficients were calculated between each cluster pair from whole and control slice datasets and visualised by a heatmap, with correlation coefficients >0.8 indicated with an asterix.

#### Differentially expressed gene analysis and gene set enrichment analysis

Differential gene expression (DEG) was determined by Welch t-tests between LMQ and control slice samples using pairwiseTTests in scran^95^ after blocking on cell type. DEG lists are detailed in Table S2. EnhancedVolcano package was used to summarise DEG results as a volcano plot.

For gene set enrichment analysis, differentially expressed genes with a p value threshold of 0.05 were selected to create a ranked gene list. Enrichment of canonical pathways gene sets (c2.cp.v2023.2.Hs.symbols.gmt) and ontology gene sets (c5.go.bp.v2023.2.Hs.symbols.gmt) from within the Human MSigDB collection were then tested by Gene Set Enrichment Analysis (GSEA) using the fgsea package^96–98^ against ranked gene lists for each cell type. Gene sets were filtered with a p value of 0.5, minimum size of 15, maximum size of 400 and number of permutations set at 10,000. Enriched gene lists and leading edge genes referred to in the text are listed in Table S3.

#### Differential Abundance Analysis

T and B cells were selectively depleted during transcriptomic analysis, so were excluded from differential abundance analyses. Differential abundance was determined by performing an ANOVA test where the overall error sum of squares and degrees of freedom are calculated from a linear model of cell type and sample type (additive model).

#### CellPhoneDB

To analyse cell-cell communication molecules, CellPhoneDB v5 (method 3) was used to understand the differentially expressed interactions. Relevant interactions were determined by: 1) interaction genes of interest must be expressed in the corresponding cell type by more than 10% of cells, and 2) at least one gene-cell type pair is identified in the DEG analysis described above. Interactions were also scored as described by CellPhoneDB v5 to rank interactions by specificity of their interacting partners^57,99^ . Results were visualised using ktplots. Relevant interactions and significant means are detailed in Table S4.

#### Packages used

Other packages used not mentioned already include ggplot2, dittoSeq, single cell experiment.

#### Multiplexed Fluorescence Imaging

Staining and imaging were performed as described^100,101^. In brief, 5μm formalin-fixed paraffin embedded (FFPE) LN slides were deparaffinised and rehydrated. Slides were then permeabilised in 0.3% Triton X-100 and washed in PBS. Antigen retrieval was performed using the NxGen decloaking chamber (Biocare Medical, Pacheco, CA, USA) for 20 min each in boiling pH6 Citrate (Agilent, S1699) and pH9 Tris-based antigen retrieval solutions. Slides were then blocked for 1 h at room temperature in PBS with 3% BSA (Merck, A7906) and 10% Donkey serum (Bio-Rad, C06SB). Slides were then washed and stained with DAPI (Thermo, D3571) for 15min, before being washed in PBS and coverslipped with mounting medial combining 50% glycerol (Sigma, G5516) and 4% propyl gallate (Sigma, 2370). The GE Cell DIVE system was used to image at 20X through iterative rounds of bleaching and staining, with slides de-coverslipped in PBS between each round. Background imaging was used to subtract autofluorescence from all subsequent rounds of staining. Staining was performed with three antibodies at a time, in blocking buffer (PBS, 3% BSA, 10% donkey serum). Bleaching was performed using NaHCO_3_ (0.1M, pH 11.2; Sigma S6297) and 3% H_2_O_2_ (Merck, 216763). The antibodies, reagents, instruments and software used are listed in Table S7. Images were analysed with QuPath v0.4.3^102^.

#### Supernatant Analysis by LEGENDPlex

At the end of culture, supernatants were collected from individual slice cultures, centrifuged to pellet contaminating cells and the supernatant transferred and stored at -80°C. Thawed supernatants were analysed for released cytokines by the LEGENDplex Human Inflammation Panel 1 (BioLegend, cat 740809) following the manufacturer’s instructions. Data was collected on an Aurora Spectral Analyzer. Cytokine concentrations were averaged from 3-4 slices per condition for each donor. Adjuvant-mediated stimulation was assessed by Wilcoxon matched-pairs signed rank test and the effect of inhibitor addition was analysed by mixed effects analysis with Dunnett’s multiple comparisons to LMQ alone.

#### Flow cytometry

Stained single cell suspensions were analysed on an LSRFortessa™ X20 (BD), and analysed by FlowJo v10.8.1 (BD Life Sciences), and LegendPlex data collected on an Aurora spectral cytometer (Cytek Biosciences) and analysed by LEGENDplex™ Data Analysis Software Suite. The antibodies, reagents, instruments, and software used are listed in Table S6.

## QUANTIFICATION AND STATISTICAL ANALYSIS

Statistical analysis was performed in GraphPad Prism v10 (GraphPad Software, LLC) or R version 4.3.2 (R Core Team (2023). R: A Language and Environment for Statistical Computing, R Foundation for Statistical Computing, Vienna, Austria. https://www.R-project.org/) and the statistical tests used are detailed in figure legends.

## SUPPLEMENTARY METHODS

### Pig Samples

Porcine mesenteric lymph nodes from healthy piglets were kindly provided by The Pirbright Institute, and were processed as for human lymph nodes.

### Quantitative real-time polymerase chain reaction (qRT-PCR)

Slices were stored in RNA protect (Qiagen) until RNA extraction by RNeasy Mini Kit (Qiagen) following manufacturer’s instructions. An electronic pestle was used to aid tissue disruption. Extracted RNA was reverse transcribed using First-strand cDNA synthesis kit (Thermo Fisher Scientific) and 100ng of complementary DNA used in a 20μl qRT-PCR reaction. PCR cycling consisted of 20 seconds at 95°C followed by 40 cycles of 3 seconds at 95°C followed by 20 seconds at 60°C. Amplified DNA products were detected by SYBR Green PCR Master Mix (ThermoFisher Scientific) and primers for IL1b, IL6 and 18S rRNA and with a ViiA 7 Real-Time PCR System, (Thermo Fisher Scientific). The reagents, instruments, and software used are listed in Table S8.

### Cell recruitment to the draining lymph node

8–12-week-old female inbred BALB/C mice (Envigo, UK) were immunised intramuscularly with a total volume of 50ul containing 25μl of LMQ in the tibialis muscle under light isoflurane anaesthesia. Draining inguinal lymph nodes were harvested at the indicated time points and digested in RPMI containing 0.8mg/ml Dispase, 0.2mg/ml Collagenase-P and 0.1mg/ml DNase, with mixing at 37°C for 3x 20 mins incubations. Cells were collected in cold PBS supplemented with 2% FCS and 2mM EDTA and filtered through a 100μM cell strainer, before being stained for flow cytometric analysis. Precision count beads™ (BioLegend) were included for calculation of absolute numbers of neutrophils (CD45+CD3−B220−Ly6C+Ly6G++), monocytes (CD45+CD3−B220−Ly6C++Ly6G+) and dendritic cells (CD45+CD3−B220−Ly6C−Ly6G−CD11c+MHCII+).

---

**Supplementary Figure 1:**
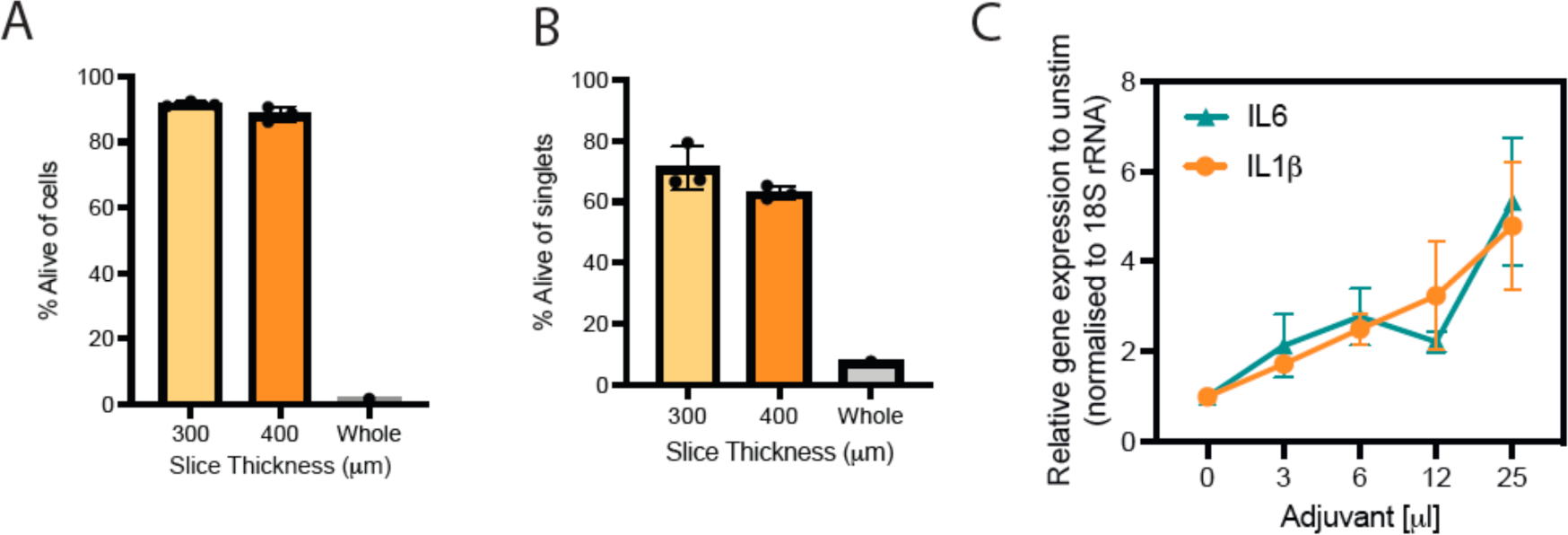
Viability and responsiveness of LN slices in culture. A) Percentage of live cells (L/D negative amongst cells gated on FSC vs SSC) from LN sections of pig LNs of indicated thickness compared to cultured whole LN after 20hr culture. (n=3 for slices, n=1 whole LN) B) Percentage of live cells (L/D negative amongst all events, gated on singlets only) from pig LN sections of indicated thickness compared to cultured whole LN after 20 h culture (n=3 for slices, n=1 whole LN) C) Expression of *IL6* and *IL1β* relative to housekeeping *18S rRNA* in pig LN slices stimulated with the indicated volumes of adjuvant in 1ml of media after 20 h of culture (n=3, average of duplicate wells)

**Supplementary Figure 2:**
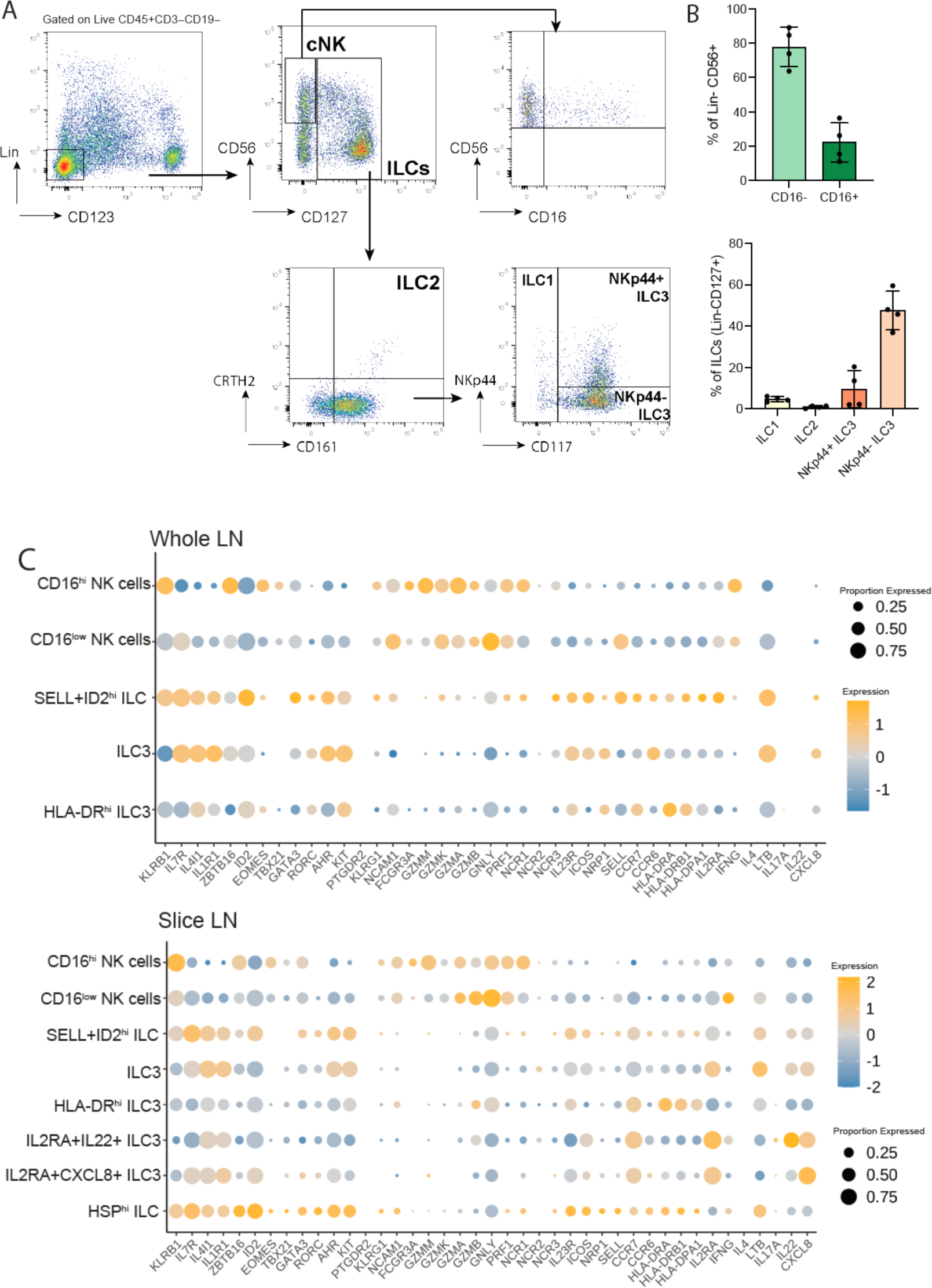
ILC populations of the human cystic LN. A) Gating strategy for the identification of NK cells and ILC subsets from whole human LN by flow cytometry. B) Quantification of the percentage of NK cells either CD16^−^ or CD16^+^ (above) and ILC subsets (below), as gated in A. C) Dot plots showing expression of marker genes for each ILC cluster within whole (above) and slice (below) human LN. Colour indicates relative log-normalised level of expression across clusters and dot size the proportion of each cluster expressing each gene.

**Supplementary Figure 3:**
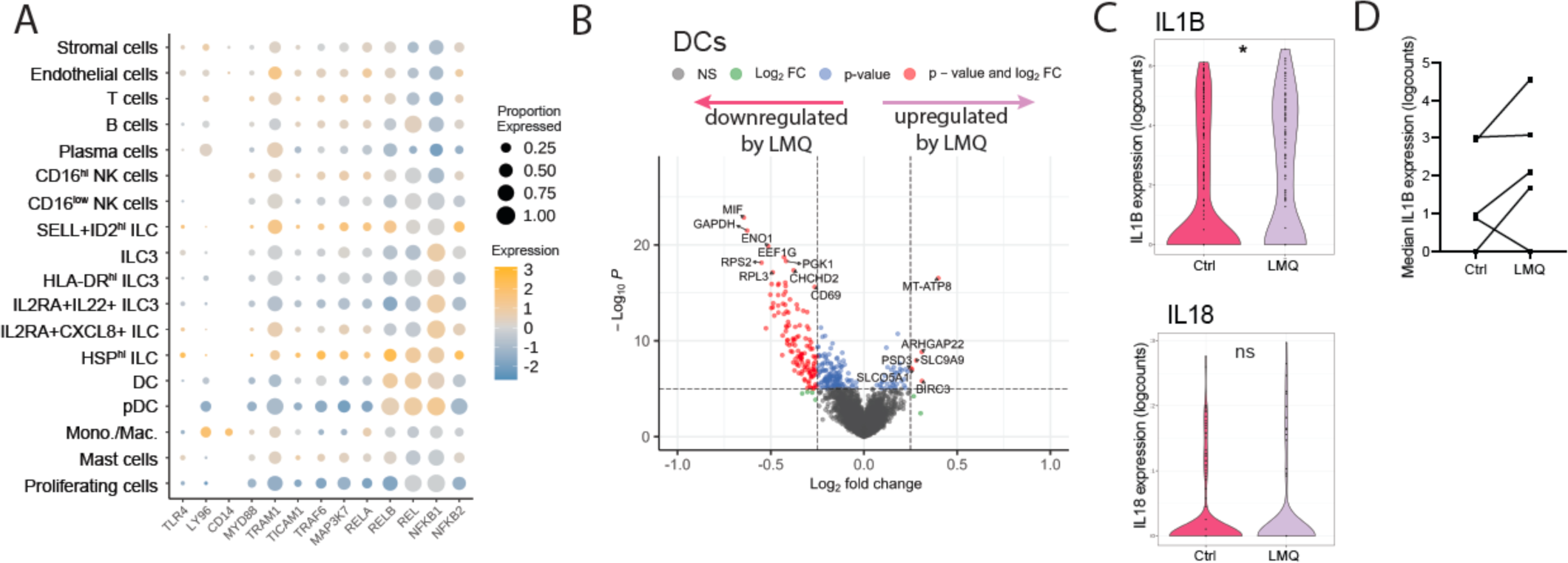
LN slice responses to TLR4 and inflammasome activation by adjuvant LMQ. A) Dot plot showing expression of TLR4 signalling components for each cell cluster within untreated LN slices. Colour indicates relative log-normalised level of expression across clusters and dot size the proportion of each cluster expressing each gene. B) Volcano plot indicating differentially expressed genes between LMQ-stimulated (right: upregulated) and control (left: upregulated) condsitions in DCs. Genes with log_2_ fold-change <0.25 are indicated in green, those with p value <10^-5^ in blue and those fulfilling both criteria in red. C) Log normalised expression levels of *IL1B* and *IL18* transcripts in the Monocytes/Macrophages cluster of all donors in control (magenta) and LMQ (lavender) treated samples. *p<0.05 D) Median log normalised expression levels of *IL1B* transcripts in the Monocytes/Macrophages cluster of paired donor slice conditions, either control or LMQ treated conditions.

**Supplementary Figure 4:**
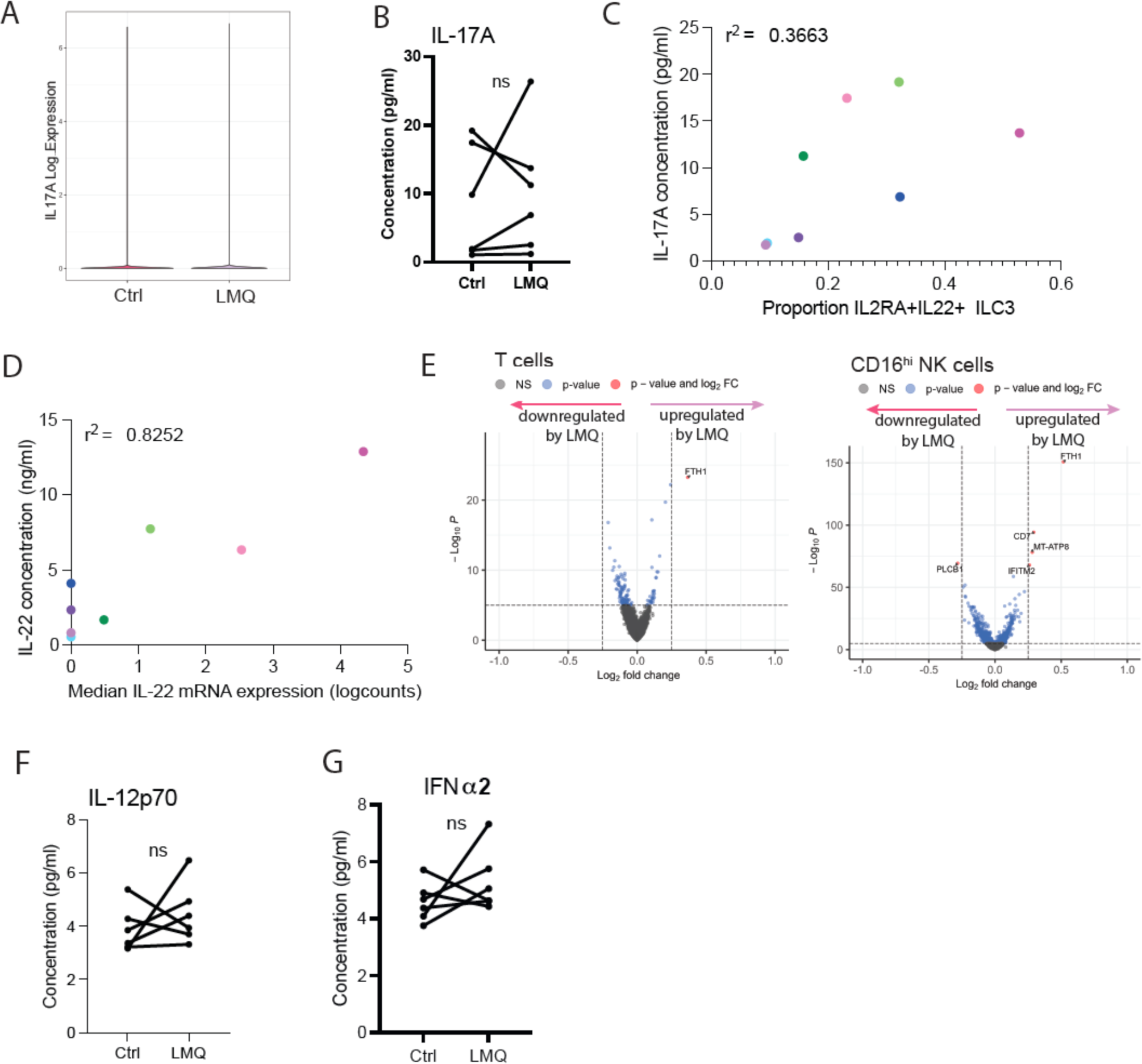
Activation of ILCs by adjuvant and downstream signalling to B cells. A) Log normalised expression levels of IL17A transcripts in all donors in control (magenta) and LMQ (lavender) treated samples. B) Cytokine concentrations of IL-17A (n=6) in slice culture supernatants after 20 h without (Ctrl) or with LMQ adjuvant (LMQ) from paired donor conditions where ns = not significant by Wilcoxon matched-pairs signed rank test. C) Proportion of IL2RA+IL22+ ILC3 cells per donor and per condition against IL-17A concentration detected in culture supernatants. Dots are coloured according to donor, and shade according to treatment (light = control, dark = LMQ treated). R^2^ value was calculated by simple linear regression. D) Median log normalised expression of IL-22 in the IL2RA+IL22+ ILC3 cluster per donor and per condition against IL-22 concentration detected in culture supernatants. Dots are coloured according to donor, and shade according to treatment (light = control, dark = LMQ treated). R^2^ value was calculated by simple linear regression. E) Volcano plot indicating differentially expressed genes between LMQ-stimulated (right: upregulated) and control (left: upregulated) conditions in T cells (left) and CD16^hi^ NK cells (right). Genes with log_2_ fold-change < 0.25 are indicated in green, those with p value <10^-5^ in blue and those fulfilling both criteria in red. F) Cytokine concentrations of IL-12p70 (n=6) in slice culture supernatants after 20 h without (Ctrl) or with LMQ adjuvant (LMQ) from paired donor conditions where ns = not significant by Wilcoxon matched-pairs signed rank test. G) Cytokine concentrations of IFNα (n=6) in slice culture supernatants after 20 h without (Ctrl) or with LMQ adjuvant (LMQ) from paired donor conditions where ns = not significant by Wilcoxon matched-pairs signed rank test.

**Supplementary Figure 5:**
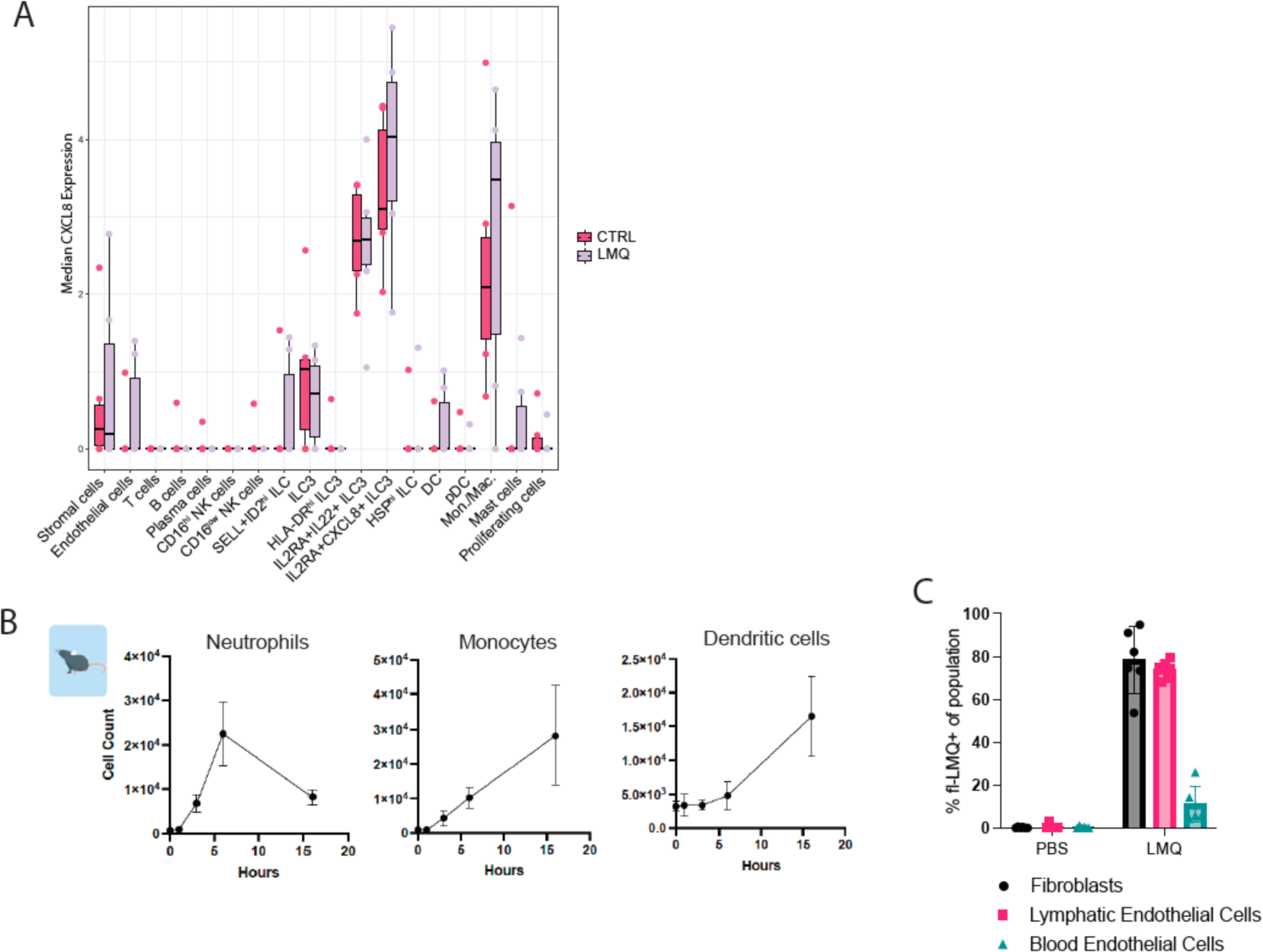
Recruitment of innate cells following i.m. injection of LMQ in mice. A) Median *CXCL8* transcript expression across cell types in human lymph node slices without (Ctrl; magenta) or with LMQ adjuvant (LMQ; lavender), each dot represents an individual donor B) Absolute cell counts of neutrophils and monocytes in the mouse draining LN at various timepoints following i.m. injection of LMQ adjuvant (n=6). C) Proportion of adjuvant positive cell populations within CD45-populations of mouse draining LN following i.m. injection of fluorescent LMQ adjuvant slices after 24h. Stromal cells were defined as CD45^−^Pdpn^+^CD31^−^ and endothelial cells as CD45^−^CD31^+^ and either Pdpn^+^ (Lymphatic Endothelial cells) or Pdpn^−^ (Blood endothelial cells). Bars indicate mean with standard deviation.

